# Transcriptome Mining Reveals a Spectrum of RNA Viruses in Primitive Plants

**DOI:** 10.1101/2022.02.07.479358

**Authors:** Jonathon C.O. Mifsud, Rachael V. Gallagher, Edward C. Holmes, Jemma L. Geoghegan

## Abstract

Current knowledge of plant viruses stems largely from those affecting economically important plants. Yet, plant species in cultivation represent a small and bias subset of the plant kingdom. Here, we describe virus diversity and abundance from a survey of 1079 transcriptomes from species across the breadth of the plant kingdom (Archaeplastida) by analysing open-source data from the One Thousand Plant Transcriptomes Initiative (1KP). We identified 104 potentially novel viruses, of which 40% comprised single-stranded positive-sense RNA viruses across eight orders, including members of the *Hepelivirales*, *Tymovirales*, *Cryppavirales*, *Martellivirales* and *Picornavirales*. One-third of the newly described viruses comprised double-stranded RNA viruses from the orders *Durnavirales* and *Ghabrivirales*. The remaining were negative-sense RNA viruses from the *Rhabdoviridae*, *Aspiviridae, Yueviridae, Phenuiviridae* and the newly proposed *Viridisbunyaviridae.* Our analysis considerably expands the known host range of 13 virus families to include lower plants (e.g., *Benyviridae* and *Secoviridae*) and four virus families to include algae hosts (e.g., *Tymoviridae* and *Chrysoviridae)*. The discovery of the first 30 kDa movement protein in a non-vascular plant, suggests that the acquisition of plant virus movement proteins occurred prior to the emergence of the plant vascular system. More broadly, however, a co-phylogeny analysis revealed that the evolutionary history of these families is largely driven by cross-species transmission events. Together, these data highlight that numerous RNA virus families are associated with older evolutionary plant lineages than previously thought and that the scarcity of RNA viruses found in lower plants to date likely reflects a lack of investigation rather than their absence.

**Importance:** Our knowledge of plant viruses is mainly limited to those infecting economically important host species. In particular, we know little about those viruses infecting primitive plant lineages such as the ferns, lycophytes, bryophytes and charophytes. To expand this understanding, we conducted a broad-scale viral survey of species across the breadth of the plant kingdom. We find that primitive plants harbour a wide diversity of RNA viruses including some that are sufficiently divergent to comprise a new virus family. The primitive plant virome we reveal offers key insights into the evolutionary history of core plant virus gene modules and genome segments. More broadly, this work emphasises that the scarcity of viruses found in these species to date likely reflects the absence of research in this area.

## 1. Introduction

Viruses are responsible for almost 50% of all emerging plant disease (1). Historically, virus identification and characterisation have focused on pathogenic viruses that infect species of economic importance with 69% of the current phytovirosphere — the total assemblage of viruses across the plant kingdom — discovered in cultivated plant species even though they represent less than 0.17% of all known plant diversity (2, 3). Importantly, the advent of metagenomic sequencing technology enables the comprehensive screening of plant tissues for novel and known viruses (4). Despite this, virus diversity in the vast majority of plants remains unquantified (5).

Our ability to infer the origins and diversification of the phytovirosphere from genomic data requires adequate sampling of the viruses across the plant kingdom. Several key plant groups are severely underrepresented or absent in previous studies of the phytovirosphere, including green algae (excluding the Chlorophytes), lower plants, gymnosperms and several angiosperm orders (5, 6). Improving knowledge across these groups will undoubtedly help uncover the evolutionary history of plant virus lineages. For instance, an analysis of the evolutionary history of viruses from algal ancestors might reveal deep associations that shaped the trajectory of plant evolution, including how the key evolutionary transitions of plants – such as terrestrialisation – have shaped the contemporary land plant virome (5). Similarly, through broad sampling across the plant kingdom, we can gain a stronger understanding of the acquisition of viruses through cross-species transmission from plant-associated organisms such as invertebrates, fungi, or protists (5).

The majority (68%) of the currently documented genera of plant viruses have positive-sense single-stranded RNA (+ssRNA) genomes and the majority of virus diversity is known only from angiosperms (7) (Figure 1). Currently, 16 viruses belonging to 12 virus families have been found in gymnosperms (8–12). Outside of several viruses found in ferns, we know little of the diversity of viruses in the lycophytes, bryophytes and charophytes that together encompass ∼27,000 species (13–16) (Figure 1). A partial analysis of published transcriptome data detected homologs of the canonical RNA virus RNA-dependent RNA polymerase (RdRp) in algae, several lower plants and gymnosperms (17). However, it is yet to be determined whether viruses that infect freshwater algae – that include the *Zygnematophyceae* ancestors of land plants – resemble those infecting angiosperms or that of the green algae (chlorophytes) which are dominated by double-stranded DNA (dsDNA) viruses particularly from the *Phycodnaviridae* (18). To date, two +ssRNA viruses related to the benyvirids have been identified in freshwater algae (19, 20). Unlike the Chlorophyta, the Charophyta characteristically contain plasmodesmata and homologs of the key components of the land plant innate immune system, both of which have been speculated to explain the absence of double-strand (ds) DNA viruses in land plants (5, 21, 22). An understanding of the viruses infecting the Charophyta and other lower plants is required to effectively test these ideas.

**Figure 1.**
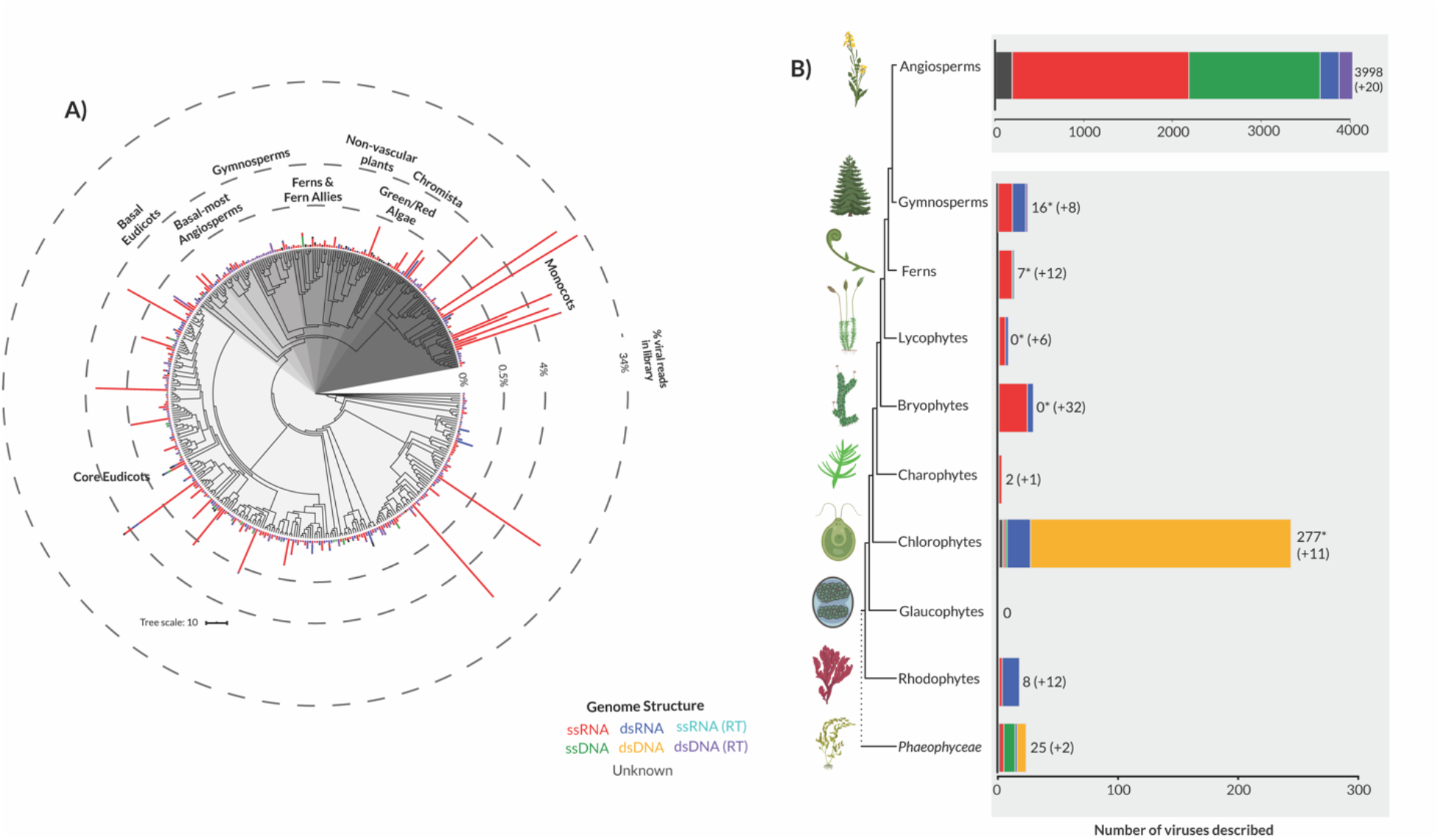
(A) Phylogram of virus composition across the One Thousand Plant Transcriptomes Initiative (1KP) samples. Plant-associated virus abundance was summarised for each plant species and normalised using a Box-Cox transformation. The height of each bar represents the percentage of virus reads detected in each plant species (after the removal of host reads). Plant clades are labelled and differentiated by shades of grey. The 1KP ASTRAL tree was used as the basis for this tree (30). Clade and abundance annotations were added using the Interactive Tree of Life (iTOL) web-based tool (109). (B) The phytovirosphere across the Plantae and *Phaeophyceae*. A schematic tree of the evolution of major plant groups. Each bar represents the number of total viruses formally or likely associated with each host group and is coloured by virus genome composition. The total number of viruses for each plant group plus those found in this study is also shown at the end of each bar. The Virus-Host (38) and NCBI virus databases (110) combined with literature searches were used to obtain virus counts. Lineage branches are not drawn to scale. To our knowledge, no viruses have been found in the Glaucophytes. Plant and algae images were obtained from BioRender.com or drawn in Adobe Illustrator (https://www.adobe.com). *Transcriptome scaffolds from libraries belonging to these host groups shared homology to virus RdRps and were partially analysed but not assembled or deposited to GenBank (17).

Transcriptome mining has become an inexpensive and efficient method of virus discovery that leverages previous investment (23–29). To this end, we mined the transcriptome data generated by the One Thousand Plant Transcriptomes Initiative (1KP) using sequence homology searches of known plant viruses. The 1KP project provides a major untapped source of polyA-selected transcriptome data for virus discovery drawn from species across the breadth of the plants in a broad sense including green plants (Viridiplantae), glaucophytes (Glaucophyta), red algae (Rhodophyta) (30, 31). Our broad aim was to revise our understanding of the phytovirosphere using data across the plant kingdom and undertake phylogenetic analyses of plant viruses to provide insights into their origins and diversification.

## 2. Methods

### 2.1 Transcriptome data generation

The 1KP generated RNA sequencing libraries from 1,143 species across the breadth of the plant kingdom (30). In addition, 30 Chromista and red alga species were also included. Due to the diversity of species examined, samples were obtained from multiple sources including field collections, greenhouses, culture collections and laboratory specimens (32). For the majority of species, young leaves or shoots were collected, although occasionally a mix of vegetative and reproductive tissues was used. To avoid RNA degradation, RNA extraction was performed immediately after tissue collection or tissue was frozen in liquid nitrogen and stored in a -80°C until extraction (32). Several extraction protocols were used including CTAB and TRIzol (see (32) for complete details). All sequencing was conducted at BGI-Shenzhen, China, using a combination of in-house protocols or TruSeq chemistry (32). All libraries were prepared from polyA RNA. Paired-end sequencing was initially completed using Illumina GAII machines (11% of libraries) with a ∼72bp read length but later the HiSeq platform was used (89% of libraries) with a 90 bp read length (32).

### 2.2 Surveying for viruses in the 1KP

Raw transcriptomes (n = 1079, belonging to 960 plants species) from the 1KP major release were downloaded from the NCBI Short Read Archive (SRA) database (BioProject accession PRJEB21674) and converted to FASTQ format using the SRA Toolkit program fastq-dump in combination with the parallel-fastq-dump wrapper (https://github.com/rvalieris/parallel-fastq-dump) (33). One hundred transcriptomes within the BioProject were not publicly available (released 22/08/2019) at the commencement of this study and thus not analysed.

Transcriptomes from the 1KP pilot study (BioProject accession PRJEB4921) and secondary project (BioProject accession PRJEB8056) were similarly not analysed. To reduce the downstream computing resources needed, raw sequences were mapped to their respective host genome scaffold using bowtie2 (34). Genome scaffolds were assembled as part of a previous study (30). Where genome scaffolds were not available (n = 2) all reads were assembled *de novo*. Trinity RNA-seq (v2.1.1) was used to quality trim and assemble *de novo* the unaligned reads captured from mapping (35). The assembled contigs were then assigned to known virus families and annotated through similarity searches against the NCBI nucleotide database (nt), the non-redundant protein database (nr) and a custom viral RdRp database using BLASTN and Diamond (BLASTX) (36, 37). To filter out weak BLAST sequence matches an e-value cut-off of 1 × 10^−10^ was employed. To identify potential false positives, putative viral contigs were manually compared across the three BLAST searches (nt, nr and RdRp) to ensure matches to virus-associated sequences were consistent.

### 2.3 Virus filtering and abundance calculations

For all analyses, we focused on virus families known to infect plants or algae. As our analyses rely on sequence-based similarity searches for virus detection it is necessarily biased towards viruses that exhibit to existing virus families. Together, the Virus-Host database (38) and the International Committee on Taxonomy of Viruses (39) were used to develop a list of plant virus families and genera to filter out virus-like contigs associated with vertebrate, invertebrate or fungi hosts based upon their top BLASTx and BLASTn matches. Packages within the Tidyverse collection (v1.3.0) in RStudio were used to complete these tasks (40–42). Where the host was ambiguous (e.g., belonged to a family or genera known to infect both plant and fungal species) the contig was inspected manually.

The relative abundance of each transcript within the host transcriptome was calculated using RNA-Seq by Expectation-Maximization (v1.2.28) (43). To account for variation in the number of unaligned reads between libraries after mapping, contig abundance was standardised by the total number of unaligned paired reads. Contigs under 200 nucleotides in length were excluded from further analysis.

### 2.4 Genome extension and annotation

Where a novel virus-like contig was discovered, we re-assembled the complete library – without removing host reads – in an attempt to recover a complete virus genome. For all re-assembled libraries, we recalculated abundance measurements to account for both host and non-host reads. The recalculated abundance measurements are shown in Supplementary Table 4. We further re-assembled all libraries belonging to non-flowering plants (n = 402). Reads were mapped onto virus-like contigs using Bbmap and heterogeneous coverage and potential misassemblies were manually resolved using Geneious (v11.0.9) (44, 45).

To determine whether a virus was novel, we followed the criteria as specified by The International Committee on Taxonomy of Viruses (39) (http://www.ictvonline.org/). Novel viruses were named using a combination of the host common name - if documented – and the associated virus taxonomic group (e.g., *Interrupted club-moss deltapartitivirus*). In cases where host assignment proved difficult the suffix “associated” was added to the host name to signify this (e.g., *Calypogeia fissa associated deltaflexivirus*). Where the taxonomic position of a virus was ambiguous the suffix “-like” was used (e.g., *Goldenrod fern qin-like virus*). Virus acronyms were created using a combination of the first and/or second letters of the host common name - if documented – and virus taxonomic group (e.g., *Leucodon julaceus beny-like virus* (LjBV)). Where multiple related viruses were found in the same host, we assigned each a number (e.g., *Odontoschisma prostratum bunyavirus 3* (OdprBV3)).

The percentage identity among virus sequences was calculated via multiple sequence alignments using Clustal Omega (v1.2.3) (46). The RdRp protein coding domain was used for all sequence alignments. Percentage identity matrices were converted to heat map plots using a custom R script provided by (28).

To characterise functional domains, predicted protein sequences along with their closest viral relatives were subjected to a domain-based search using the Conserved Domain Database (v3.18) (https://www.ncbi.nlm.nih.gov/Structure/cdd/cdd.shtml) and cross-referenced with the PFAM (v34.0) and Uniclust30 (v2018_08) databases available within the MMseqs2 webserver (47). To recover additional annotations, we used HHpred within the MPI Bioinformatics Toolkit webserver to query the PDB_mmCIF70 (v.12_Oct), SCOPe70 (v2.08), UniProt-SwissProt-viral70 (v3_Nov_2021) and TIGRFAMs (v15.0) databases (48). Virus genome diagrams were produced using the program littlegenomes (49). Where available NCBI/GenBank CDS information was used to annotate reference virus sequences (50).

### 2.5 Detection of endogenous virus elements

All genome scaffolds produced by the 1KP were used as a database in which we queried using the protein translations of the viruses discovered in this study. Endogenous viral elements (i.e., EVEs) were detected using the tblastn algorithm (51). The search threshold was limited to 100 amino acids in length with an e-value cut off of 1×10^−20^. Where multiple hits across several plant scaffolds were observed we manually examined the sequence. Suspected endogenous virus sequences were queried against a subset whole-genome shotgun contig database which included green plants (taxid: 33090) and red algae (taxid: 2763). In addition, the virus-like sequences discovered in this study were checked for host gene contamination using the contamination function implemented in CheckV (v0.8.1) (52). All potential endogenous sequences were removed from further analyses.

### 2.6 Assessing library contamination by eukaryotes, bacteria, and protozoa

For libraries in which a novel virus was discovered we investigated whether reads belonging to other eukaryotes were also present in the sequencing libraries. To achieve this, we obtained taxonomic identification for raw reads in each library – without the removal of host reads – by aligning them to the NCBI nt database using the KMA aligner and the CCMetagen program (53, 54). Sequence abundance was calculated by counting the number of nucleotides matching the reference sequence with an additional correction for template length (the default parameter in KMA). Krona charts generated by CCMetagen were edited were further edited in Adobe Illustrator (https://www.adobe.com) (55). Library contamination was also assessed by the 1KP and used to inform our host-virus assignments (31).

### 2.7 Phylogenetic analysis of plant viruses

Phylogenetic trees of the plant-associated viruses discovered here were inferred using a maximum likelihood approach. We combined our translated virus contigs with known virus protein sequences from each respective virus family taken from NCBI/GenBank (50). Sequences were then aligned with the program Clustal Omega (v1.2.3) with default parameters (46). Sites of ambiguity were removed using trimAl (v1.2) (56). To estimate phylogenetic trees, selection of the best-fit model of amino acid substitution was determined using the Akaike information criterion, corrected AIC, and the Bayesian information criterion with the ModelFinder function (-m MFP) in IQ-TREE (57, 58). All phylogenetic trees were created using IQ-TREE with 1000 bootstrap replicates. Phylogenetic trees were annotated with FigTree (v1.4.4) (59) and further edited in Adobe Illustrator (https://www.adobe.com).

To visualise the occurrence of cross-species transmission and virus-host co-divergence across plant virus families, we reconciled the co-phylogenetic relationship between viruses and their hosts. For each select plant virus family, a vascular plant host cladogram was constructed using trees from (60) and (61), using the R package V.PhyloMaker (v0.1.0) (62). As lower plants and non-plant species are not present in the V.PhyloMaker megatree, these hosts were added to the cladogram using the software phyloT, a phylogenetic tree generator based on NCBI taxonomy (http://phylot.biobyte.de/) as well as topologies available in the appropriate literature. The host information was obtained from the NCBI Virus database (accessed 14/12/2021) and available literature (63) A tanglegram that graphically represents the correspondence between host and virus trees was created using the R packages phytools (v0.7-80) and APE (v5.5) (64, 65). Virus sequences from each family were obtained through a broad survey of all virus genomic data available on GenBank. The virus phylogenies used in the co-phylogenies were constructed as detailed above. To quantify the relative frequencies of cross-species transmission versus virus-host co-divergence we reconciled the co-phylogenetic relationship between viruses and their hosts using the Jane co-phylogenetic software package (66). Jane employs a maximum parsimony approach to determine the best ‘map’ of the virus phylogeny onto the host phylogeny. The cost of duplication, host-jump and extinction event types were set to one, while host-virus co-divergence was set to zero as it was considered the likely null event. Following the parsimony principle, the reconciliation proceeds by minimising the total event cost. The number of generations and the population size was both set to 100. Jane was chosen over its successor eMPRess as it allows for a virus to be associated with multiple host species and handle polytomies (67). For a multi-host virus, we represented each association as a polytomy on the virus phylogeny.

### 2.8 Assigning plant host clades

Each plant host was assigned to each clade in a previous study based upon their phylogenetic positioning and lineage information (30). To improve clarity when colouring the phylogenies (although not the tanglegrams) we reduced the number of clades from 25 to ten (core eudicots, basal eudicots, monocots, basalmost angiosperms, gymnosperms, fern and fern allies, non-vascular, green algae, red algae and lastly Chromista) by combining those that were closely related or potentially overlapping to increase the number of species in each group (SI Table 1).

### 2.9 Data availability

The raw One Thousand Plant Transcriptomes Initiative sequence reads are available at BioProject PRJEB21674. All viral genomes and corresponding sequences assembled in this study have been deposited in NCBI GenBank and assigned accession numbers xxxx-yyyy.

## 3. Results

We characterised the viruses found in the transcriptomes of 960 plant species within the 1KP major release. The transcriptomes represented a broad taxonomic sampling across the Archaeplastida (green plants, glaucophytes and red algae). Sequencing libraries had a median of 25,187,714 paired reads (range 10,156,464–46,650,336). A median of 82% of reads (range 1%-96%) in these libraries mapped to host genome scaffolds and were subsequently removed. *De novo* assembly of the sequencing reads resulted in a median of 36,015 contigs (range 1,396–146,217) per library, with a total of 41,256,176 contigs generated (SI Table 2).

### 3.1 Diversity and abundance of plant viruses

In total, virus-like transcripts were found for 603 plant species; 69% of these were plant-associated while numerous identified sequences shared high similarity to non-plant associated viruses including those known to infect fungi, invertebrate and vertebrate hosts. Among the non-plant-associated virus transcripts, 34% were unclassified (10% of total virus-like transcripts) such that they were most closely related to a virus sequence with little to no taxonomic information (i.e., a virus sequence classified as only belonging to the *Riboviria*). If an RdRp-like region was detected in an unclassified virus-like transcript we further assessed whether it could be plant-associated (see Phylogenetic analysis of identified viruses). The remaining non-plant-associated virus transcripts were largely classified within the *Orthomyxoviridae* (vertebrate associated) (25%), *Rhabdoviridae* (invertebrate associated) (17%), *Partitiviridae* (fungus associated) (10%), *Mimiviridae* (amoeboid associated) (10%) and *Adenoviridae* (vertebrate associated) (7%) and excluded from the remainder of this study. These sequences are discussed in more detail in the section on “Presence of contaminants in sequencing libraries” below. Although some of these viruses could represent plant infection it remains challenging to discern and we, therefore, made the conservative decision to remove them from the analysis.

We detected transcripts closely associated with viruses containing single and double-stranded DNA and RNA genomes. The majority of virus-like sequences belonged to families with +ssRNA genomes (61%) or reverse-transcribing dsDNA viruses (22%) (Figure 1). The +ssRNA virus transcripts were predominately classified within the *Betaflexiviridae* (30%), *Potyviridae* (19%), *Secoviridae* (16%) and *Alphaflexiviridae* (10%) (SI Table 3). Negative-sense single-stranded RNA (-ssRNA) virus transcripts were classified within the *Aspiviridae* (0.04%), *Rhabdoviridae* (6%) and *Tospoviridae* (3%) (*Phenuiviridae* and *Yueviridae* transcripts were later detected in the unclassified virus-like transcripts) (SI Table 3). dsDNA virus transcripts with sequence similarities to the *Phycodnaviridae* were detected across the algae samples. These phycodna-like virus transcripts frequently encoded the chitinase and DNA ligase genes which are homologous to those in distantly related host organisms including fungi and bacteria. Due to the difficulties discerning whether these transcripts represent *Phycodnaviridae* sequences or contamination, we excluded all phycodnavirus*-*related sequences. All remaining dsDNA viruses were exclusively reverse-transcribing viruses from the *Caulimoviridae*. We failed to detect any sequences that shared homology with several plant virus families including *Reoviridae*, *Nanoviridae* and *Fimoviridae* (although see the Discussion for caveats).

There was a large range of total viral abundance in each library (5×10^-6^% – 31% reads after host-associated reads were removed). Viruses with +ssRNA genomes accounted for the vast majority (99.8%) of virus abundance detected (Figure 1, SI Table 3). As expected, virus discovery was concentrated in the flowering plants (angiosperms), which have the highest number of previously classified viruses. For instance, plant virus-like sequences were frequently discovered in the core eudicots and monocots (i.e., 73% of libraries in which plant virus transcripts were found). The detection rate of plant viruses was highest in the most basal angiosperms (57%) and monocots (50%). No significant difference in virus abundance was observed between sequencing platforms (Genome Analyzer II and Illumina HiSeq 2000; p=0.327).

### 3.2 Presence of contaminants in sequencing libraries

The bacterial, fungal and insect species that live in or on plant tissues are commonly sampled within plant sequencing libraries (31), although contamination from other plants is also a possibility during sample preparation or sequencing. To quantify the extent of library contamination we used the KMA and CCMetagen tools (Figure 2). Among the libraries analysed (n = 95), bacteria were consistently detected representing a median of 1.5% of total abundance (range 0.01%-33%). A median of 2% of library abundance was associated with fungi sequences (range 0%-53%). Arthropods and chordates were also commonly detected across libraries (found in 87 and 89 libraries, respectively) but at lower abundance (median 0.15%, range 0%-11.4%). The presence of chordate associated reads is likely attributed to various routes of sample contamination (e.g., faeces) or during sample processing and sequencing.

**Figure 2.**
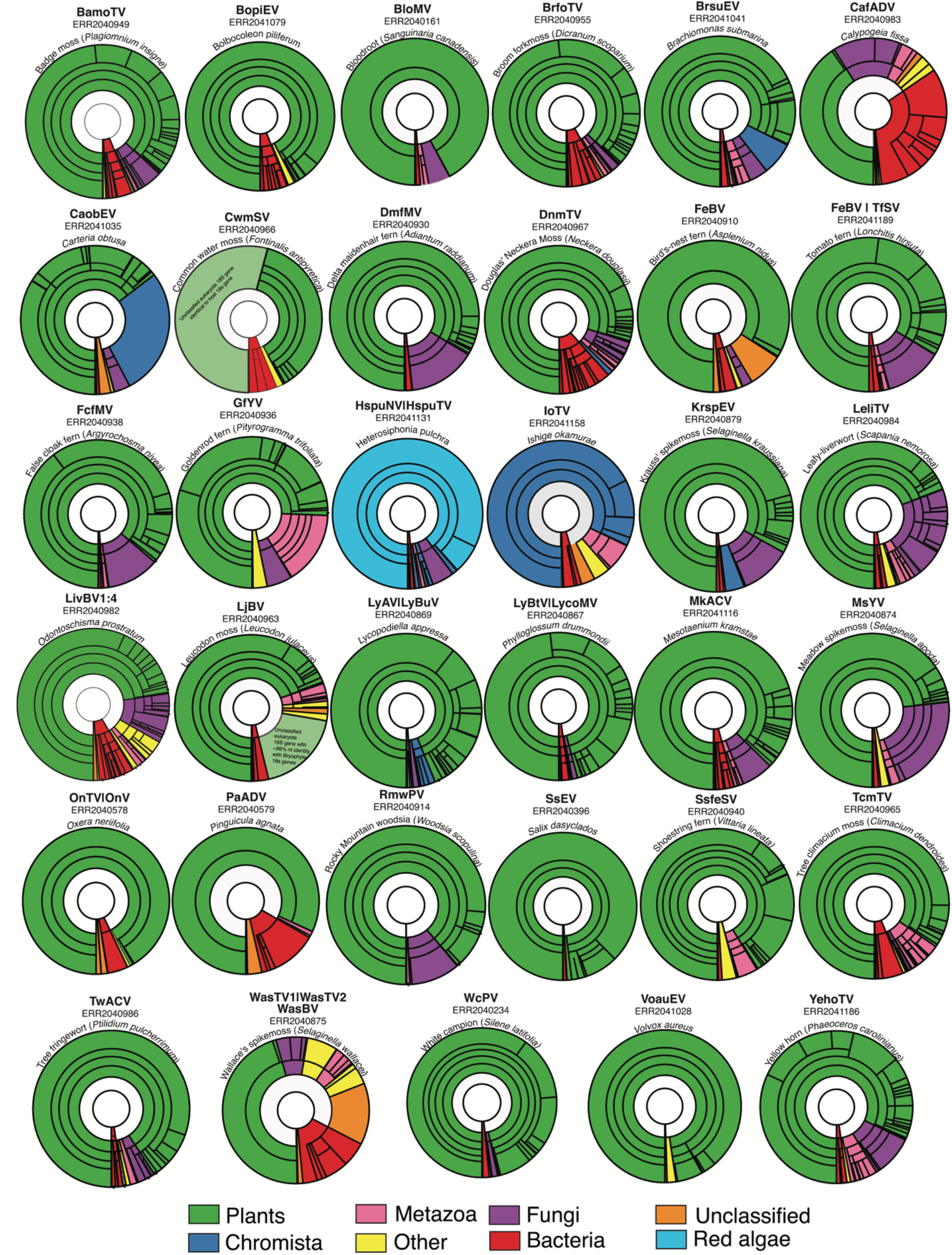
Taxonomic assignments of reads in select One Thousand Plant Transcriptomes Initiative (1KP) libraries. Each Krona graph illustrates the relative abundance of taxa in a metatranscriptome at varying taxonomic levels. For clarity, a maximum depth of five taxonomic levels was chosen for each graph. The library Sequence Read Archive accession number, host species, and the corresponding virus of interest are annotated above each graph. Segments are highlighted based upon the species taxonomic grouping (plants = green, Chromista = blue, unclassified = orange, bacteria = red, metazoa = pink, fungi = purple, red algae = light blue, other = yellow). Here “plants” encompasses the Viridiplantae. Reads without any match in the nt database are not shown.

The detection of four vertebrate associated viruses across several libraries provided further evidence of library contamination. Sequences belonging to these viruses - *Influenza A virus* (16 libraries), *Human mastadenovirus C* (30 libraries), *Human immunodeficiency virus* (15 libraries) and *Parainfluenza virus 5* (3 libraries) – were present at low abundance and showed little genetic variation between libraries. Notably, chordate-associated reads were only present in 66% of libraries in which these viruses were found. The failure to consistently detect potential hosts for these viruses suggests contamination during sequencing. The four vertebrate associated viruses were largely absent in libraries in which novel plant-associated viruses were discovered, except for the *Larix speciosa*, *Brachiomonas submarina, Climacium dendroides*, *Silene latifolia and Oxera neriifolia* transcriptomes.

In addition, the 1KP compared all assembled sequences to a reference set of nuclear 18S ribosomal RNA sequences from the SILVA small subunit rRNA database using BLASTn (31, 68). Where a sample had several alignments to any other plant sequences outside of the expected source family the sample was described as having “worrisome contamination” (31). This applied to eleven plant libraries in which novel viruses were identified. Below, we discuss library contaminates from viewpoint of virus-host associations.

### 3.3 Phylogenetic analysis of identified viruses

To infer phylogenetic relationships between identified viruses, order and family-level phylogenetic trees were estimated using the highly conserved viral region that comprises the RdRp. In total, we assembled 104 RdRp contigs that likely represent novel virus species, of which 41 were considered as unclassified or non-plant associated due to their similarities to virus groups known to infect non-plant hosts (SI Table 4). Further analysis of these contigs revealed that they are likely plant-associated.

#### 3.3.1 Positive-sense single-stranded RNA ((+)ssRNA) viruses

##### Hepelivirales

###### Benyviridae

We identified three beny-like sequences that to our knowledge represent the first benyvirid found in lower plants. The first sequence, tentatively named *Fern benyvirus* (FeBV), was found in both the bird’s-nest fern (*Asplenium nidus*) and tomato fern (*Lonchitis hirsuta*). Together with *Wheat stripe mosaic virus*, FeBV represents a well-supported clade separate from the remaining plant benyviruses (Figure 3).

**Figure 3.**
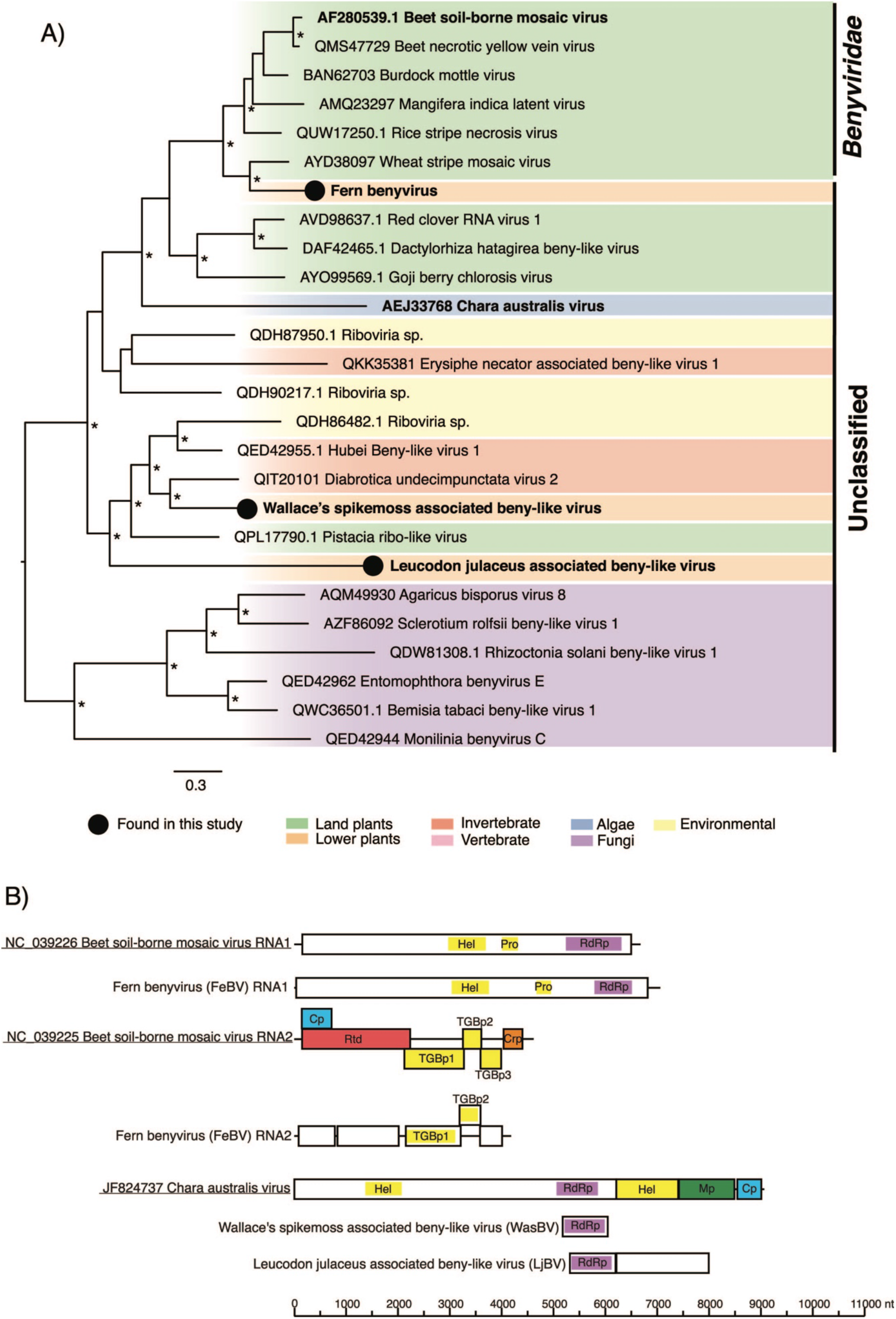
(A) Phylogenetic relationships of the beny-like viruses identified in this study. ML phylogenetic tree based on the RNA-1 replicase protein shows the topological position of virus-like sequences discovered in this study (black circles) in the context of their closest relatives. Branches are highlighted to represent host clade (land plants = green, lower plants = orange, invertebrate = red, vertebrate = pink, algae = blue, fungi = purple, yellow = environmental, Chromista = light blue, red algae = dark green). Here “Land plants” encompasses both angiosperms and gymnosperms while “Lower plants” includes the bryophytes, lycophytes, and ferns. All branches are scaled to the number of amino acid substitutions per site and trees were mid-point rooted for clarity only. An asterisk indicates node support of >70% bootstrap support. Tip labels are bolded when the genome structure is shown on the right. (B) Genomic organization of the beny-like virus sequences identified in this study and representative species used in the phylogeny. Beet soil-borne mosaic virus RNA three and four are not pictured here. The data underlying this figure and definitions of acronyms used are presented in SI Table 5.

The triple gene block (TGB) is a hallmark gene module of the *Benyviridae* among several other virus families in the class *Alsuviricetes* (69)). In both fern libraries, proteins resembling the TGB were assembled (Figure 3). The TGB proteins shared ∼34% amino acid identity with the TGB protein of other benyvirids. To our knowledge, this is the first TGB protein found outside of flowering plants. Phylogenetic analysis placed the TGB1 protein of FeBV basal to the *Benyviridae* (SI figure 1).

Two additional beny-like viruses, named here *Leucodon julaceus associated beny-like virus* (LjBV) and *Wallace’s spikemoss associated beny-like virus* (WasBV) were assembled. LjBV and WasBV cluster with unclassified algae, invertebrates, fungi and soil-derived viruses forming a group basal to all plant benyvirids and potentially constitute a novel virus group (Figure 3). LjBV contains a second open reading frame (ORF) with no detectable homology to known sequences (Figure 3).

Due to the phylogenetic placement of LjBV and WasBV close to viruses infecting distant hosts (e.g., invertebrates and fungi), we investigated the potential of contamination from other eukaryotes as the source of these viruses. Of note, the Wallace’s spikemoss metatranscriptome contained reads that matched various fungi orders (7% of all reads) as well as those matching the plant-parasitic oomycete *Albugo laibachii* (7%) which makes inferring virus-host relationships challenging (Figure 2). Reads belonging to various fungi species accounted for 10% of the bird’s-nest fern transcriptome and 12% of the tomato fern transcriptome (Figure 2). Despite the presence of fungi-associated reads, the phylogenetic position of FeBV suggests that FeBV is likely plant-associated (Figure 3). No concerning contaminants were detected in the *Leucodon julaceus* transcriptome.

##### Tymovirales

###### Betaflexiviridae

We identified 18 virus sequences that fell within the order *Tymovirales*. Four virus transcripts were associated with the *Betaflexiviridae*. The first, named *Sea beet betaflexivirus* (SbBV) clusters with *Agapanthus virus A*, an unclassified betaflexivirus (Figure 4). The remaining sequences denoted *Iranian poppy betaflexivirus* (IpBV), *Linum macraei betaflexivirus* (LimBV) and *Lycopod associated betaflexivirus* (LyBtV) resemble capilloviruses. Notably, LyBtV may extend the known host range of the *Betaflexiviridae* from angiosperms to lower plants. All sequences phylogenetically cluster with known capilloviruses and potentially represent novel virus species (Figure 4). The *Phylloglossum drummondii* library in which LyBtV was assembled had contamination from lycopod and dicot species (Figure 2). As the majority of plant-associated reads were assigned to lycophytes (50%), LyBtV has been tentatively assigned to this group.

**Figure 4.**
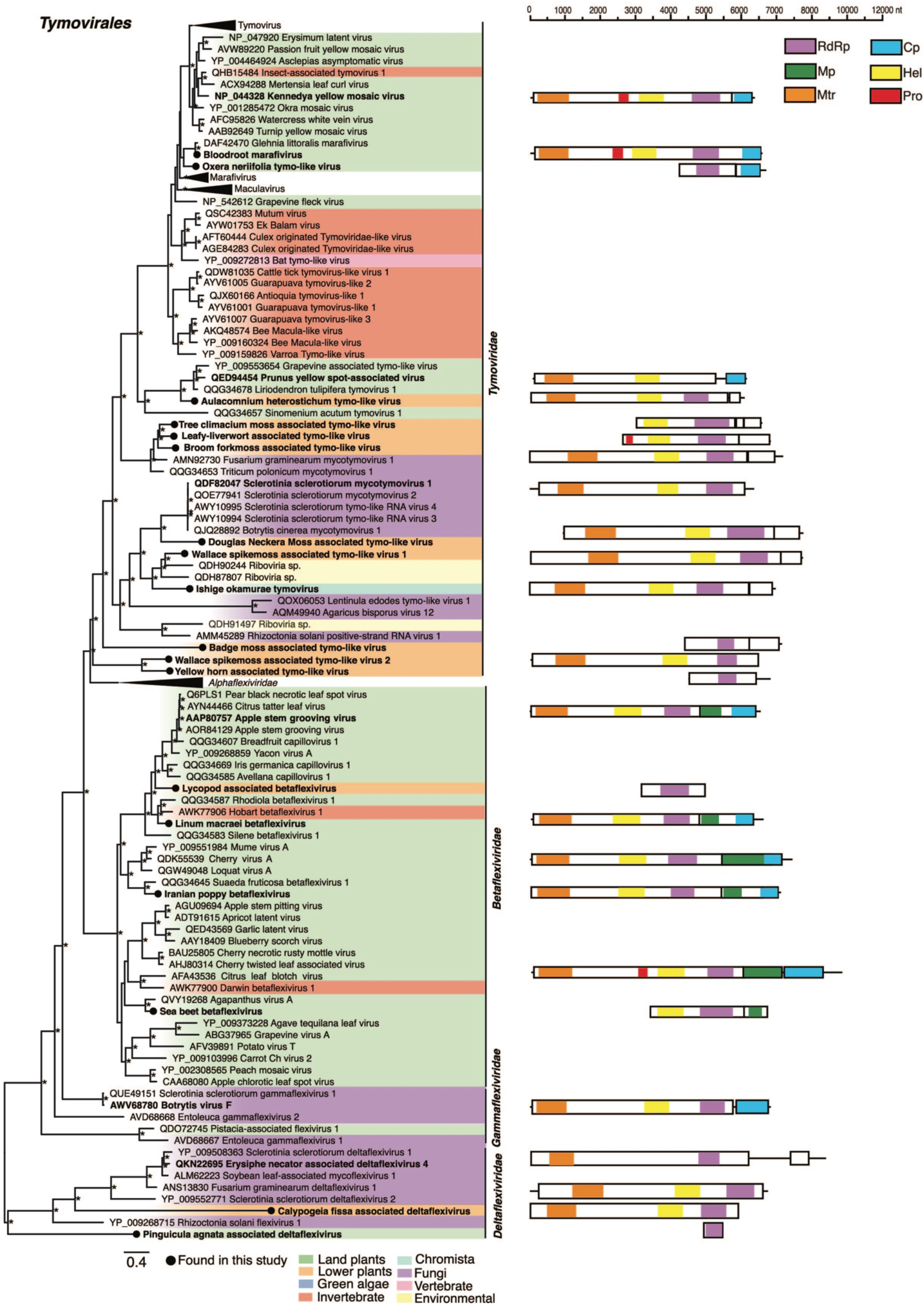
Left: Phylogenetic relationships of the viruses within the order *Tymovirales*. ML phylogenetic tree based on the replication protein shows the topological position of virus-like sequences discovered in this study (black circles) in the context of their closest relatives. See Figure 3 for the colour scheme. All branches are scaled to the number of amino acid substitutions per site and trees were mid-point rooted for clarity only. An asterisk indicates node support of >70% bootstrap support. Tip labels are bolded when the genome structure is shown on the right. Right: Genomic organization of the virus sequences identified in this study and representative species used in the phylogeny.

###### Tymoviridae

We identified 12 virus-like sequences that clustered within the *Tymoviridae* and related viruses. *Ishige okamurae associated tymo-like virus* (IoTV) was detected in the brown alga *Ishige okamurae* and likely represents the first virus in the order *Tymovirales* from brown algae. IoTV, along with ten sequences assembled from hornworts, liverworts and bryophytes grouped with tymo-like viruses from fungus and environmental samples (Figure 4). It is uncertain whether the true hosts of the novel tymo-like viruses discovered here are plants. Fungi contaminates were detected across these libraries but varied in abundance (range 1%-21%, mean = 6%). Despite their clustering with mycotymoviruses, *Broom forkmoss associated tymo-like virus* (BrfoTV) and *Tree climacium moss associated tymo-like virus* (TcmTV) were assembled from libraries with ∼1% fungal reads, highlighting the inherent difficulties in host-virus assignment. Importantly, <1% of reads in *Ishige okamurae* transcriptome belonged to species of fungi (Figure 2).

We assembled two tymo-like virus sequences denoted *Oxera neriifolia tymo-like virus* (OnTV) and *Bloodroot marafivirus* (BloMV). BloMV and OnTV grouped with the unclassified *Glehnia littoralis marafivirus* (Figure 4). Marafiviruses and tymoviruses are commonly distinguished from each other based upon a highly conserved 16 nucleotide (nt) sequence known as the “tymobox” [GAGUCUGAAUUGCUUC] in tymoviruses and the “marafibox” [CA(G/A)GGUGAAUUGCUUC] in marafiviruses (70, 71). While these two novel viruses cluster together phylogenetically, they differ in terms of genome structure and motifs. A “marafibox” like sequence appears to be present in BloMV (CAACGCGAAUUGCUUU) (5606-5621 nt) albeit differing by several residues. This finding, combined with the BloMV genome likely consisting of a single large ORF, supports the assignment of BloMV as a *Marafivirus*. OnTV, like members of the *Tymovirus* genera, contains both a second ORF – likely encoding a coat protein (CP) – and a tymobox (1493-1508 nt) (Figure 4). Phylogenetic analysis of the coat protein sequence places OnTV and BloMV in a clade with macula- and marafi-like viruses (SI Figure 1).

###### Deltaflexiviridae

We assembled two sequences that share similarities to members of the mycotymovirus family, *Deltaflexiviridae*. The first sequence was detected in the liverwort *Calypogeia fissa*, tentatively named *Calypogeia fissa associated deltaflexivirus* (CafADV) and appeared distantly related to delta- and gammaflexiviruses. A second related partial sequence, named here *Pinguicula agnata virus* (PaV), shared 32% amino acid identity with mycoflexivirus, *Botrytis virus F*. In a phylogenetic analysis with members of the *Tymovirales,* CafADV and PaAGV are placed with the deltaflexivirids (Figure 4).

It is unclear whether the source of these virus sequences is from plants or contamination from other eukaryotes. The *C. fissa* library contained numerous contaminates including algae, fungi and bacteria representing 1%, 15% and 33% of total reads respectively, which make discerning the host association for CafAV challenging (Figure 2). Interestingly, no fungi-associated reads were found in the *P. agnata* library suggesting a potential plant origin (Figure 2).

##### Picornavirales

###### Secoviridae

We identified four sequences that shared similarities to members of the *Secoviridae* denoted *Common water moss secovirus* (CwmSV), *Salix dasyclados secovirus* (SadSV), *Tomato fern secovirus* (TfSV) and *Shostring fern secovirus* (SfSV). CwmSV, TfSV and SfSV cluster within the nepoviruses and likely represented the first seco-like virus detected in the bryophytes and ferns (Figure 5).

**Figure 5.**
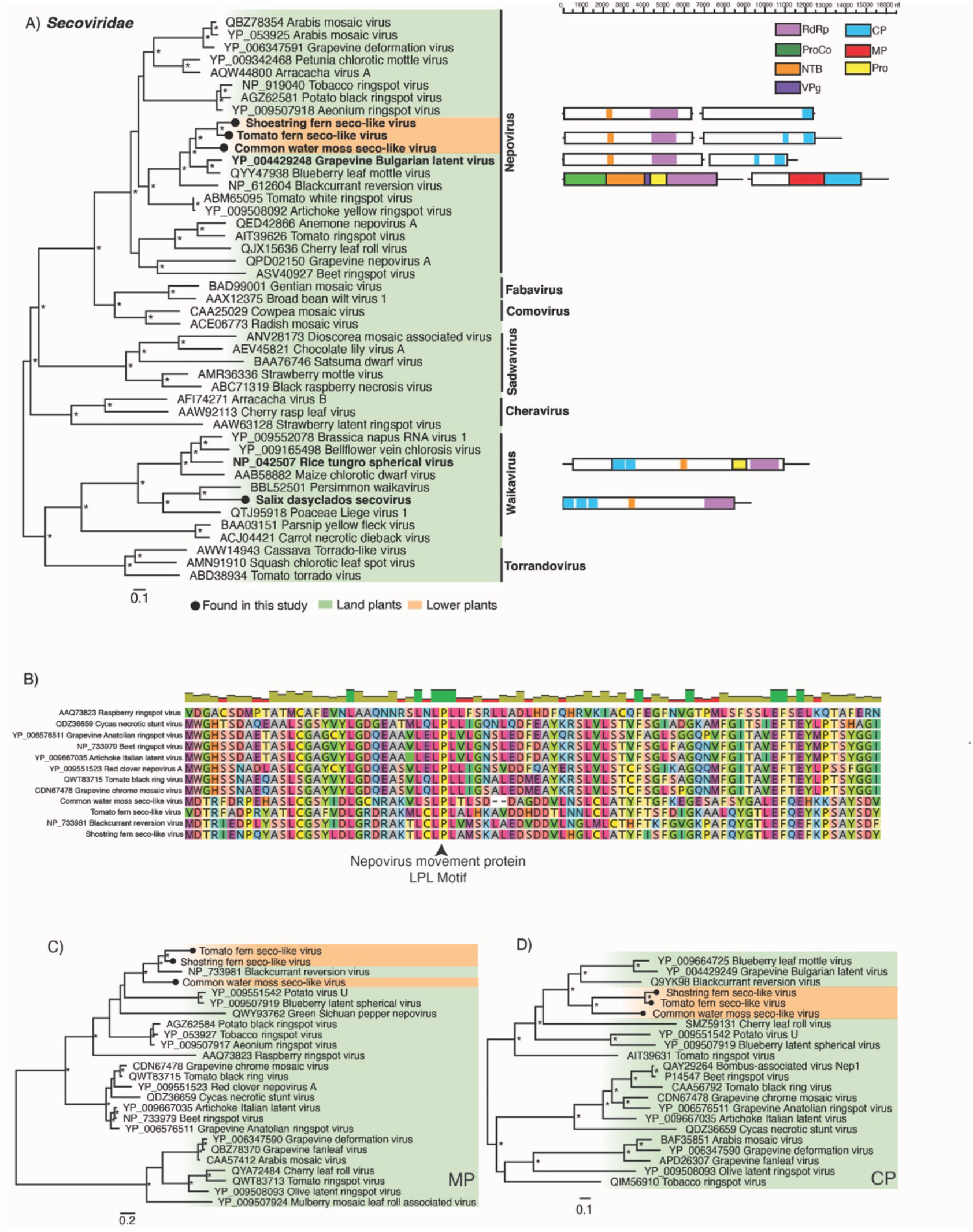
(A) Left: Phylogenetic relationships of the viruses identified within the virus family *Secoviridae*. ML phylogenetic trees based on the Pro-pol region show the topological position of virus-like sequences discovered in this study (black circles) in the context of their closest relatives. Right: Genomic organization of the seco-like sequences identified in this study and representative species used in the phylogeny. (B) Multiple amino acid sequence alignment of the 30K movement protein “LPL” motifs which are highly conserved throughout the nepoviruses. (C) Phylogenetic relationships of the Nepovirus 30K movement proteins. D) Phylogenetic relationships of the Nepovirus coat proteins. For all trees, branches are scaled to the number of amino acid substitutions per site and trees were mid-point rooted for clarity only. An asterisk indicates node support of >70% bootstrap support. Tip labels are bolded when the genome structure is shown on the right. See Figure 3 for the colour scheme. Viruses discovered in this study are signified using a black circle on the tree tip.

A putative RNA2 ORF was assembled for the three nepovirus-like sequences each containing a complete CP (Figure 5). The CPs fall within the nepovirus subgroup C (Figure 5D). While a movement protein (MP) domain was not formally detected, we predict that the region upstream of the CP contains a putative movement-like protein. For CwmSV, this region (amino acid position 312-883) displayed sequence homology to the MP of *Blackcurrant reversion virus* (E-value: 5.42e-86, amino acid identity: 46%). Both TfSV and SfSV displayed similar levels of homology in this region. We detected the LPL motif which is commonly found in nepovirus MPs in all three viruses (Figure 5B). Phylogenetic analysis of the putative MPs placed these viruses with *Blackcurrant reversion virus* in the genera Nepovirus (Figure 5C).

We found little evidence that these viruses were detected due to contamination by land plants or other eukaryotes. The *F. antipyretica* transcriptome was composed of reads closely related to a feather moss belonging to the order Hypnales to which *F. antipyretica* is also found. Furthermore, a large proportion of reads were assigned to an uncultured eukaryote 18S rRNA gene (54%) (HG421124.1) that was identical to the *F. antipyretica* 18S rRNA (AF023714.1) among other bryophyte 18S rRNA genes in a blastn search (e-value = 2e-102, nucleotide identity = 100%) (Figure 2). Fungi represented 12% of reads in the *L. hirsute* transcriptome. Despite this, it is unlikely that TfSV is fungi-associated as no fungal contamination was detected in the *Vittaria lineata* transcriptome in which the closely related SfSV sequence (amino acid identity: 78%) was assembled (Figure 2).

##### Lenarviricota

###### Mitoviridae

We identified six virus sequences that cluster within the *Mitoviridae* - denoted *Chinese swamp cypress mitovirus* (CscMV)*, Asian bayberry mitovirus* (AsbaMV)*, False cloak ferns mitovirus* (FcfMV)*, Delta maidenhair fern mitovirus* (DmfMV) and *Lycopod associated mitovirus* (LycoMV). The fern (FcfMV and DmfMV) and lycophyte (LycoMV) associated sequences cluster with the fern *Azolla filiculoides mitovirus 1* and form a sister group to the plant mitoviruses and non-retroviral endogenous RNA viral elements (NERVEs) (Figure 6) (16). The gymnosperm associated sequences form a sister all the plant-associated mitoviruses and NERVEs. KpcMV extends the known host range of plant mitoviruses from ferns to lycophytes. Another mito-virus sequence was detected in the green alga *Bolbocoleon piliferum*, denoted *Bolbocoleon piliferum mito-like virus* (BopiMV). BopiMV falls basal to the mitoviruses, distinct from various unclassified mito-like viruses including the green algae associated *mito-like picolinusvirus* (QOW97241) (Figure 6). All novel sequences show strong conservation of the motifs characteristic of mitovirus RdRps (SI Figure 2) (72).

**Figure 6.**
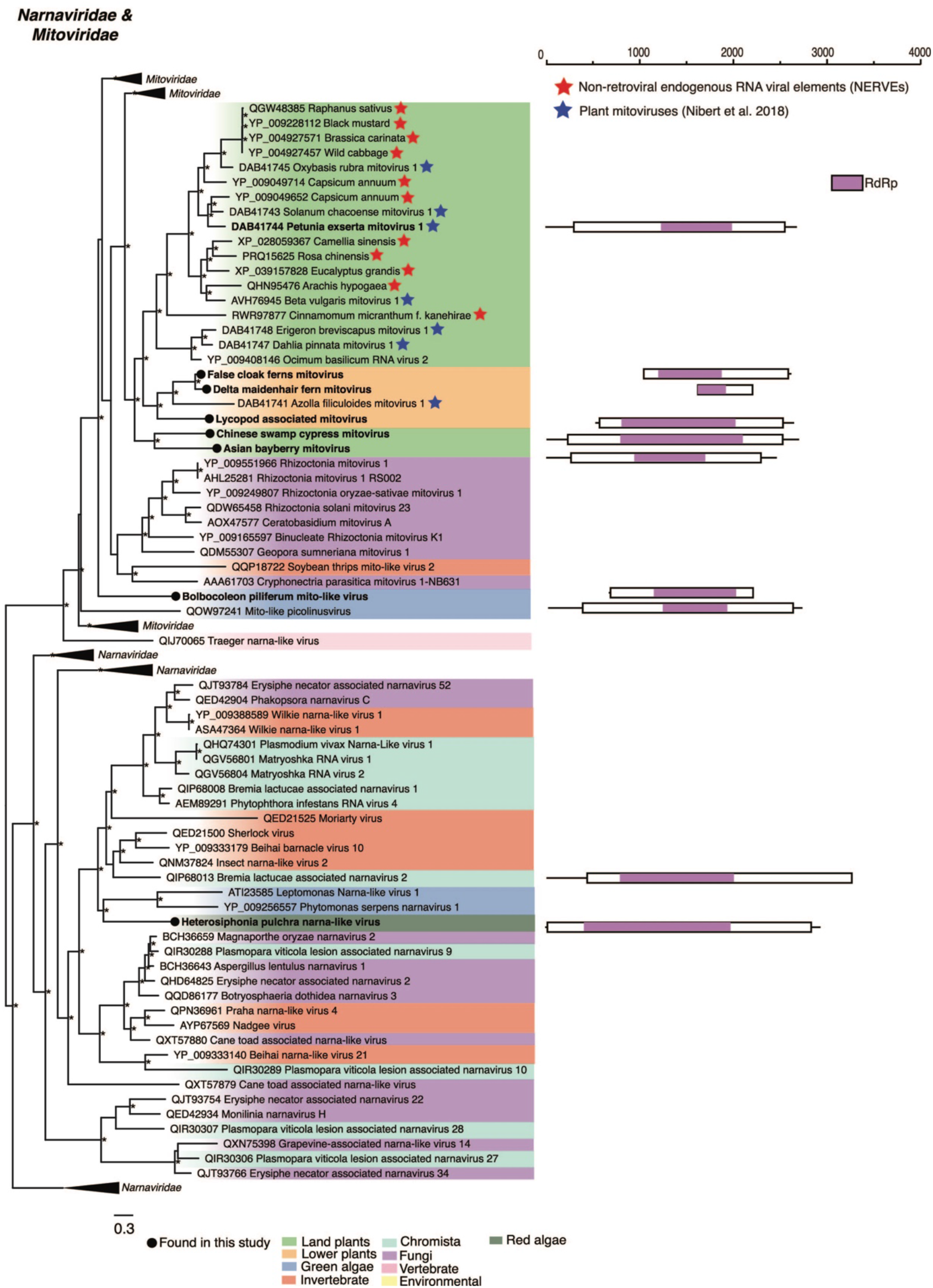
Left: Phylogenetic relationships of the viruses within the virus families *Narnaviridae* and *Mitoviridae*. ML phylogenetic trees based on the replication protein show the topological position of virus-like sequences discovered in this study (black circles) in the context of their closest relatives. See Figure 3 for the colour scheme. Blue stars signify mitovirus sequences identified in (16). Red stars signify non-retroviral endogenous RNA viral elements (NERVEs). All branches are scaled to the number of amino acid substitutions per site and trees were mid-point rooted for clarity only. An asterisk indicates node support of >70% bootstrap support. Tip labels are bolded when the genome structure is shown on the right. Right: Genomic organization of the virus sequences identified in this study and representative species used in the phylogeny.

There is little evidence to suggest that these sequences are derived from a non-plant organism. While the FcfMV and DmfMV libraries were contaminated with fungi, (12% and 15% of reads respectively) fungi-associated reads were absent in the libraries of all other mitoviruses. As the codon UGA encodes tryptophan (Trp) in fungal mitochondria this codon assignment is also present in fungal mitoviruses (73–75). In contrast, the UGA codon in plant mitochondria is a stop codon and hence absent from plant mitovirus sequences except as a stop codon (16). The absence of internal UGA codons in these sequences is further evidence that these sequences are plant-derived (16, 76). Although additional analyses are required, we found no evidence through searches of the 1KP genome scaffolds and the WGS shotgun database that these sequences are mitochondrial or nuclear NERVEs. Furthermore, CscMV, AsbaMV and LycoMV contain complete RdRps and their UTRs share similarities in length and identity with plant mitoviruses.

###### Narnaviridae

A partial narna-like virus sequence was identified in the red alga *Heterosiphonia pulchra* denoted Heterosiphonia pulchra narna-like virus (HspuNV). HspuNV clusters with unclassified trypanosomatid associated viruses. While ∼5% of reads in this library were associated with fungi the phylogenetic position of this virus suggests that it is not derived from fungi (Figure 2, Figure 6).

##### Tolivirales

###### Tombusviridae

An alphacarmo-like virus tentatively named *Ihi tombusvirus* (IhiTV) was identified in an Ihi (*Portulaca molokiniensis*) sample. IhiTV is phylogenetically positioned within the alphacarmoviruses (SI Figure 3).

##### Patatavirales

###### Potyviridae

We identified three virus-like sequences that clustered with plant viruses in the family *Potyviridae* – *Traubia modesta potyvirus* (TramPV), *Common milkweed potyvirus* (ComPV) and *Salt wort potyvirus* (SawPV). TramPV and ComPV shared 87% amino acid identity and may therefore represent a single virus species. The potyvirus-like sequences discovered all group with known potyviruses in a phylogenetic analysis of the Nib gene (SI Figure 3).

##### Martellivirales

###### Endornaviridae

Six alphaendorna-like virus sequences were detected in the four green algae species and one lycophyte. The green algae and lycophyte associated alphaendorna-like viruses termed *Bolbocoleon piliferum endorna-like virus* (BopiEV), *Volvox aureus endorna-like virus* (VoauEV), *Carteria obtusa associated endorna-like virus* (CaobEV), *Brachiomonas submarina associated endorna-like virus* (BrsuEV), *Staurastrum sebaldi endornavirus* (SsEV) and *Krauss’ spikemoss associated endorna-like virus* (KrspEV) fall across the alphaendornavirus phylogeny and predominately cluster with algae and fungi associated viruses (Figure 7). There was little evidence of algae (non-host) or fungi contamination in the *S. sebaldi*, *B. piliferum* and *V. aureus* transcriptomes with <1% of all reads associated with these groups (Figure 2). Non-green algae contaminants were present in the *C. obtus* (28%)*, B. submarina* (7%) and *S. kraussiana* (4%) transcriptomes where fungi also appeared as a notable contaminate representing 11% of all reads (Figure 2). To our knowledge, these sequences represent the first endornavirus associated with charophytes, chlorophytes and lycophytes although further work is needed to confirm the virus-host associations.

**Figure 7.**
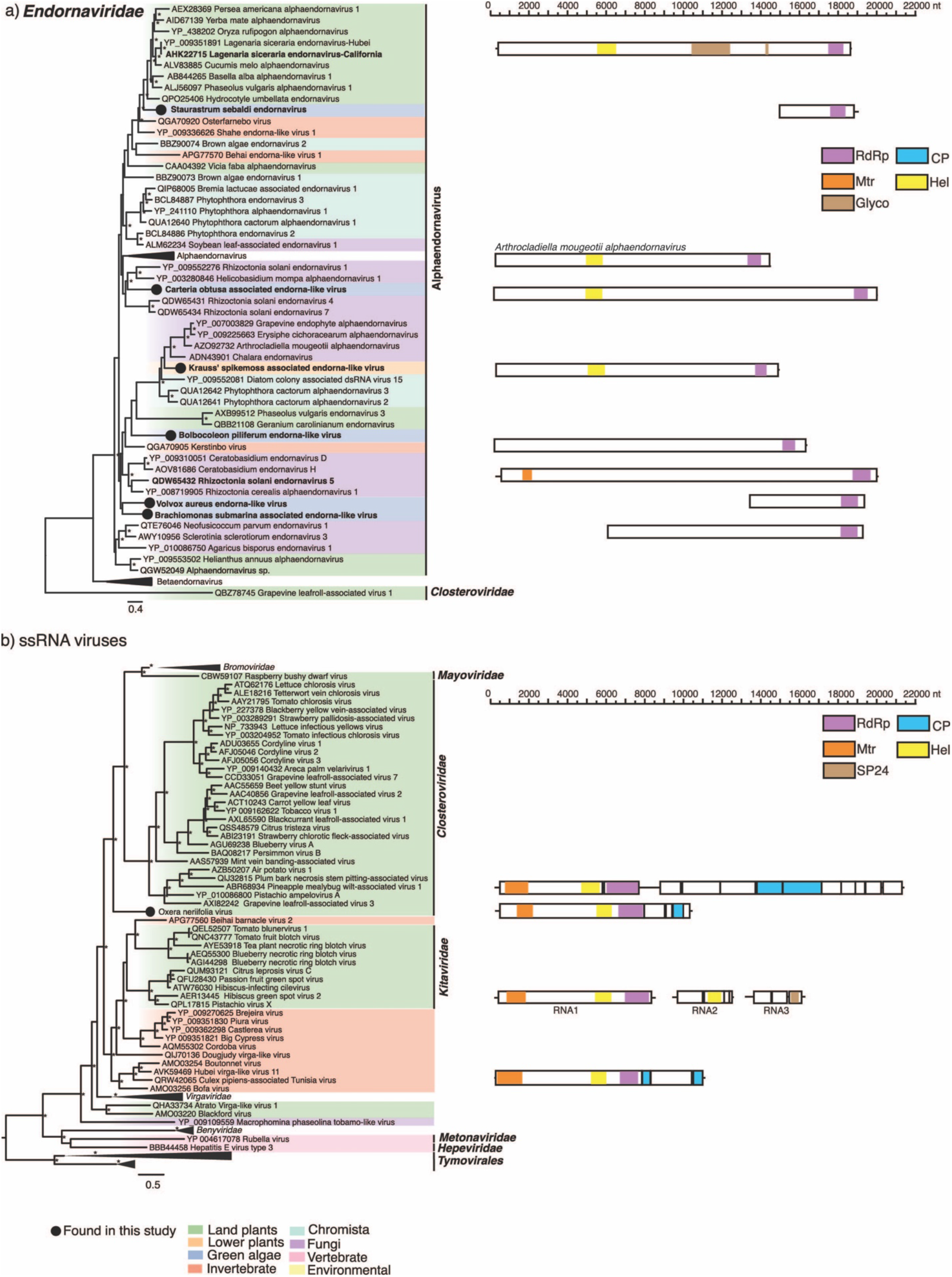
Left: Phylogenetic relationships of the (A) endorna-like and (B) unclassified (+)ssRNA virus identified in this study. ML phylogenetic trees based on the replication protein show the topological position of virus-like sequences discovered in this study in the context of those obtained previously. Right: Genomic organization of the (A) endorna-like and (B) unclassified ssRNA virus sequence identified in this study and representative species used in the phylogeny. For all trees, branches are scaled to the number of amino acid substitutions per site and trees were mid-point rooted for clarity only. An asterisk indicates node support of >70% bootstrap support. Tip labels are bolded when the genome structure is shown on the right. See Figure 3 for the colour scheme. Viruses discovered in this study are signified using a black circle on the tree tip.

###### Unclassified

We identified a virus-like sequence in an *Oxera neriifolia* library, termed *Oxera neriifolia associated virus*. The sequence, 10,214 nt in length contained four ORFs. The first ORF (7,536 nt) comprised of a viral methyltransferase, helicase, and RNA polymerase while the third ORF (513 nt) most closely resembled a CP. ORF one and ORF three shared the greatest sequence similarity with *Culex pipiens associated Tunisia virus* (32% amino acid identity). The second and fourth ORF share no homology to known viruses. The genome organization of OnV is distinct from the other related plant virus families (Figure 7). OnV forms a distinct and well-supported outgroup to the *Closterviridae, Bromoviridae* and *Mayoviridae* families. As such, OnV may potentially constitute a new virus family (Figure 7). We found little evidence that OnV was detected due to contamination by other eukaryotes (Figure 2).

#### 3.3.2 Negative-sense single-stranded RNA ((-)ssRNA) viruses

##### Bunyavirales

###### Phenuiviridae

A phenui-like virus sequence termed *Brown algae phenui-like virus* (BralPV) was recovered from a *Sargassum thunbergii* transcriptome. The partial L segment clusters with the unclassified plant and fungi viruses (Figure 8). No additional phenui-like virus segments were recovered. There were no concerning contaminants were detected in the *S. thunbergii* transcriptome (Figure 2).

**Figure 8.**
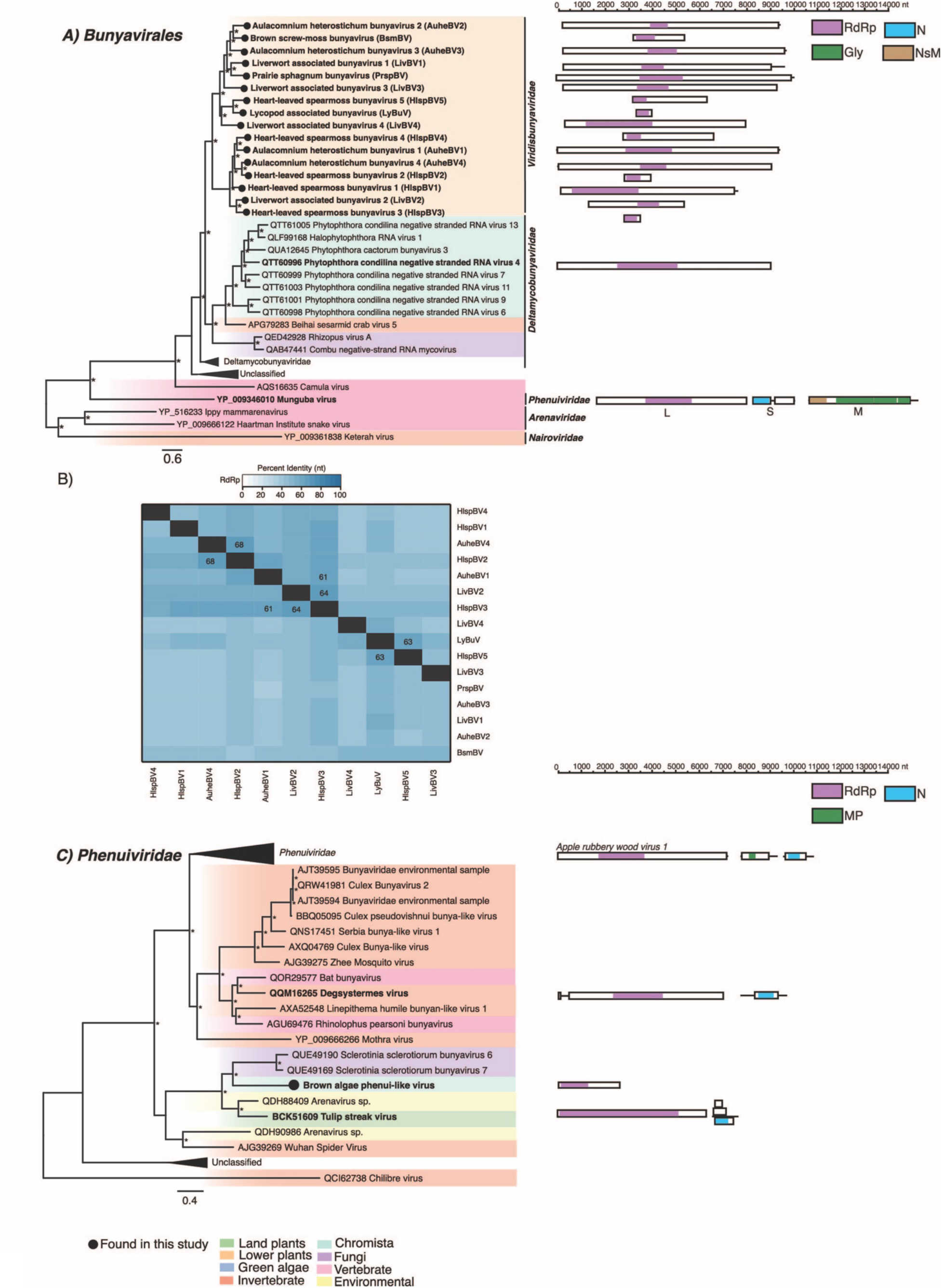
Phylogenetic relationships of the viruses (A) Left: A phylogeny depicting a novel clade of viruses related to the *Deltamycobunyaviridae* in the context of the *Bunyavirales*. Right: Genomic organization of the virus sequences identified in this study and representative species used in the phylogeny (B) Percent identity matrix of the novel bunya-like viruses. Identity scores are calculated from an alignment of the RdRp protein coding sequence. For clarity, the 100% identity along the diagonal has been removed. Where sequence identity is >= 60% the value is shown. (C) Left: A phylogeny depicting the phenui-like virus identified in this study in the context of the *Phenuiviridae*. Right: Genomic organization of the virus sequences identified in this study and representative species used in the phylogeny. For all trees, branches are scaled to the number of amino acid substitutions per site and trees were mid-point rooted for clarity only. An asterisk indicates node support of >70% bootstrap support. Tip labels are bolded when the genome structure is shown on the right. See Figure 3 for the colour scheme. Viruses discovered in this study are signified using a black circle on the tree tip.

###### Viridisbunyaviridae

We identified 16 bunya-like virus sequences from eight liverwort, moss and lycophyte libraries. Three libraries contained multiple distinct putative complete and partial viruses. The overall pairwise nucleotide identity was <70% between each sequence (Figure 8). As such we consider each a different bunya-like viruses. These sequences group together to form a novel clade of unclassified bunya-like viruses distantly related to oomycete, fungi, and invertebrate viruses (Figure 8). Bunyaviruses typically comprise three segments (L, M, and S), although only the L segment was recovered for these sequences. These sequences represent the first plant-associated viruses that cluster near the unofficially named *Deltamycobunyaviridae* (77) (Figure 8). As the complete coding sequences of the viruses discovered share <30% amino acid identity to the nearest relatives in the *Deltamycobunyaviridae,* they may constitute a new virus family. We tentatively name this virus family the *Viridisbunyaviridae*, *(Viridis* meaning green, while bunya is derived from the virus *order Bunyavirales* in which this clade falls within). There was no evidence suggesting that these sequences originated from non-plant contaminants. Host assignment was unclear for *Lycopod associated bunyavirus* and *Liverwort associated bunyavirus 1:4* as reads belonging to several lycophyte and liverwort species, respectively, were found in the source transcriptomes (Figure 2).

##### Mononegavirales

###### Rhabdoviridae

We identified seven sequences that clustered with plant viruses in the family *Rhabdoviridae* denoted *Canadian violet rhabdovirus 1* (CvRV1), Canadian violet rhabdovirus 2 (CvRV2), *Common ivy rhabdovirus* (CoiRV) and *Indian pipe rhabdovirus* (InpRV), *Tree fern varicosa-like virus* (TfVV), *Monoclea gottschei varicosa-like virus* (MgVV) and *Bug moss associated rhabdo-like virus* (BmRV). Notably, TfVV and MgVV expand the host range of the rhabdoviruses from angiosperms and gymnosperm to ferns and liverworts. RNA2 segments were recovered for both viruses, TfVV RNA2 contained five genes while MgVV contained four (Figure 9C). Two partial segments sharing similarities to the nucleocapsid (N) of *Black grass varicosavirus-like virus* (YP_009130620.1) were found in the Indian pipe library and share 50% amino acid identity. All sequences likely represent novel species within known plant infecting genera (Figure 9C).

**Figure 9.**
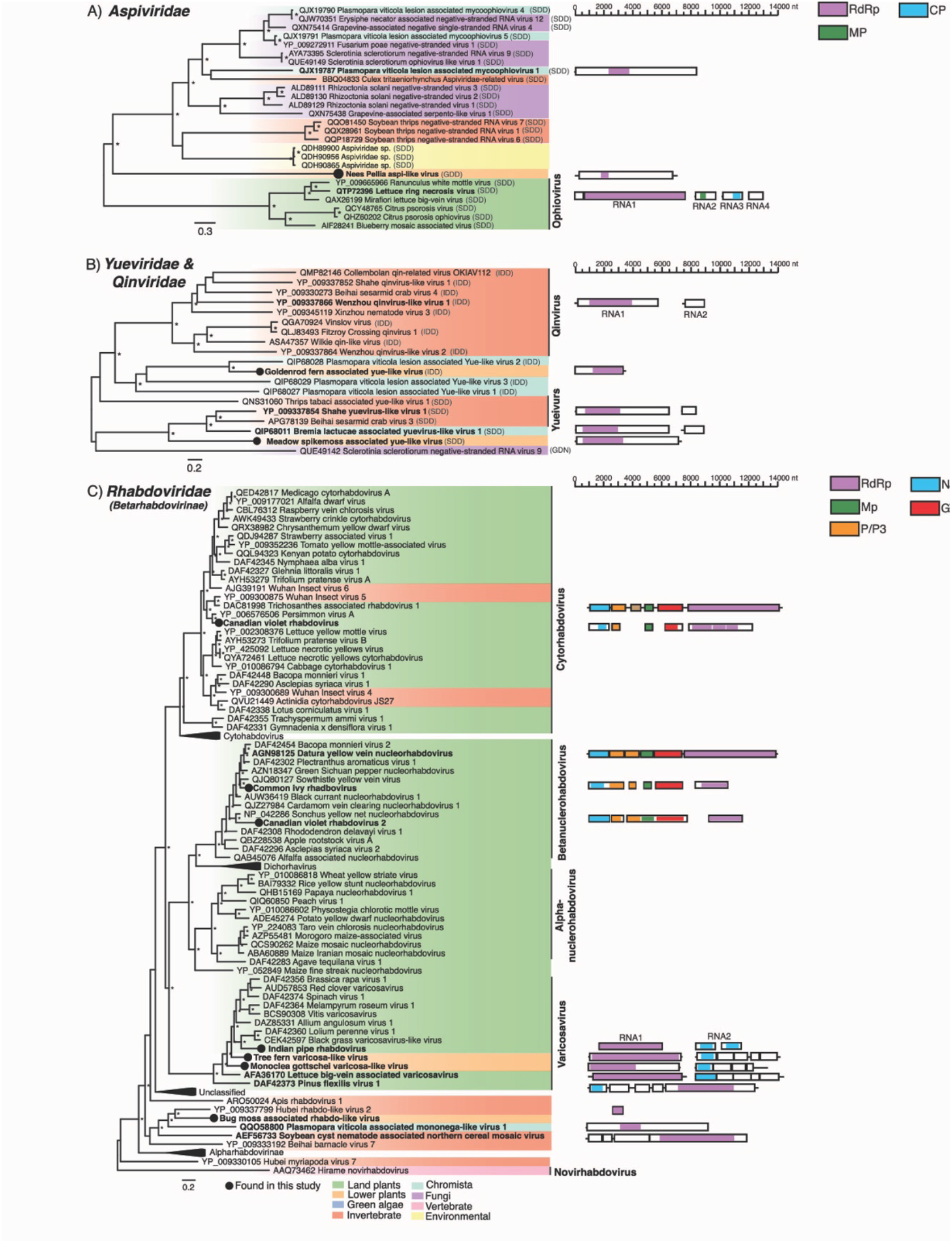
Left: Phylogenetic relationships of the viruses within the families, (A) *Aspiviridae*, (B) *Yue*- and *Qinviridae* and (C) *Rhabdoviridae.* Right: Genomic organization of the virus sequences identified in this study and representative species used in the phylogeny. For all trees, branches are scaled to the number of amino acid substitutions per site and trees were mid-point rooted for clarity only. An asterisk indicates node support of >70% bootstrap support. Tip labels are bolded when the genome structure is shown on the right. See Figure 3 for the colour scheme. Viruses discovered in this study are signified using a black circle on the tree tip. For trees (A) and (B), the RdRp motif C trimer of each sequence is shown in brackets at the end of the tip label.

BmRV is a partial sequence (693 nt) most closely related to the unclassified *Hubei rhabdo-like virus 2* (44% amino acid identity). Further evidence is needed to confirm BmRV as the first moss rhabdovirus, but the relatively low proportion of contaminates in this library (3% algae and 3% fungi) suggests that this virus is plant-associated (Figure 2). While 53% of reads in the MgVV library were fungi associated the phylogenetic position of MgVV suggests it is derived from plants (Figure 2, Figure 9C).

##### Serpentovirales

###### Aspiviridae

We identified an aspi-like sequence termed *Nees’ Pellia aspi-like virus* (NpAV). A complete RNA1 segment (6989 nt) was assembled, although no other segments were recovered (Figure 9A). NpAV most closely resembles *Rhizoctonia solani negative-stranded virus 3* (amino acid identity: 22%) and falls basal to all the unclassified aspi-like viruses including those found in fungi, invertebrates, and oomycetes. NpAV is the first aspi-like virus identified in plants outside of the angiosperms and may constitute a novel virus group (Figure 9A). Notably, unlike the other aspiviruses that possess a SDD sequence in motif C of the RdRp – a known signature for segmented negative-stranded RNA viruses – NpAV has a GDD sequence (Figure 9A).

##### Goujianvirales

###### Yueviridae

An yue-like virus sequence termed *Meadow spikemoss associated yue-like virus* (MsYV) was found in the lycophyte *Selaginella apoda* and most closely resembles algae associated *Bremia lactucae associated yuevirus-like virus 1* (amino acid identity: 26%). Phylogenetic analysis supports the assignment of MsYV as the first plant yuevirus (Figure 9B)

A second partial yue-like virus sequence was detected in a *Pityrogramma trifoliata* library and termed *Goldenrod fern associated yue-like virus* (GfYV). GfYV falls with a group of oomycete associated viruses. Consistent with the qin-like viruses, GfYV has an IDD (Ile-Asp-Asp) sequence motif instead of the common GDD (Gly-Asp-Asp) in the catalytic core of its RdRp, while MsYV contains SDD (Ser-Asp-Asp) in the same manner as many yue-like viruses (Figure 9B). The libraries from which GfYV and MsYV were assembled are contaminated with fungal reads (5% and 21%, respectively) as such host assignment is made with caution (Figure 2). Reads belonging to oomycetes were not found in either library.

#### 3.3.3 Double-stranded RNA (dsRNA) viruses

##### Durnavirales

###### Amalgaviridae

We detected five sequences that cluster with amalga-like viruses. *Lycopod associated amalgavirus* (LycoAV) is a partial RdRp containing a sequence that falls basal to the *Amalgaviridae* and represents the first amalga-like virus in the lycophytes (Figure 10). Three amalga-like sequences were discovered in green and red algae transcriptomes and cluster with *Diatom colony associated dsRNA virus 2* (Figure 10). As noted in the case of *Bryopsis mitochondria-associated dsRNA virus* and several green algae associated viruses (78) when translated into amino acids using the protozoan mitochondrial code, two overlapping ORFs are present: the first, encoding a hypothetical protein, while the second, a replicase through a −1 ribosomal frameshift (79). For two of the amalga-like sequences identified in this study – *Nucleotaenium eifelense virus* (NueiV) and *Rhodella violacea virus* (RhviV) – a similar structure was observed but we were unable to identify any ribosomal frameshift motifs in either sequence (Figure 10). Further work is needed to confirm if these sequences should be translated through the mitochondrial genetic code.

**Figure 10.**
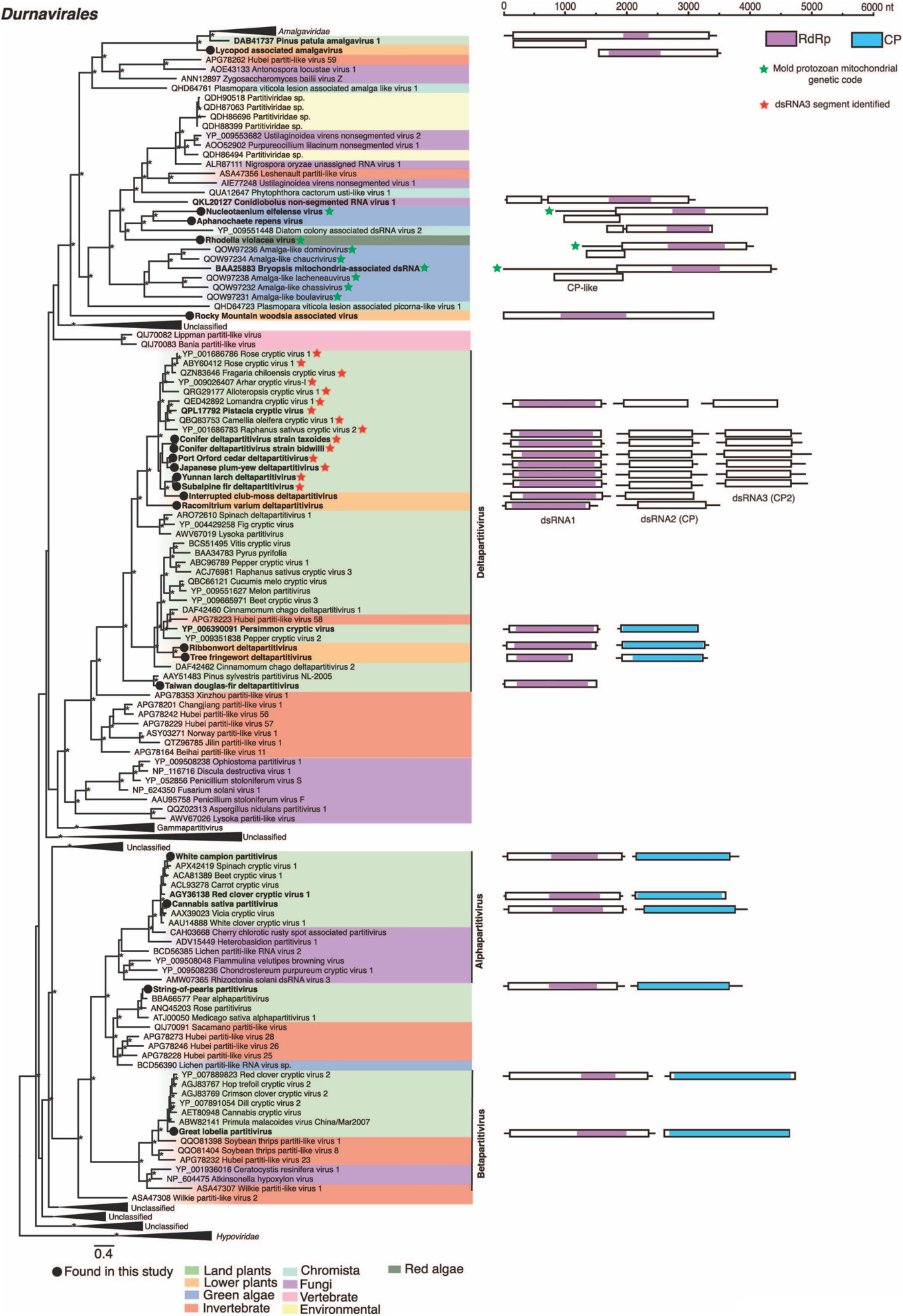
Left: Phylogenetic relationships of the viruses within the order *Durnavirales*. ML phylogenetic trees based on the replication protein show the topological position of the virus-like sequences discovered in this study (black circles) in the context of their closest relatives. See Figure 3 for the colour scheme. Green stars are used to signify sequences that have been translated using the protozoan mitochondrial genetic code. Red stars are used to signify sequences for which a dsRNA3 coat protein-like segment has been described. All branches are scaled to the number of amino acid substitutions per site and trees were mid-point rooted for clarity only. An asterisk indicates node support of >70% bootstrap support. Tip labels are bolded when the genome structure is shown on the right. Right: Genomic organization of the virus sequences identified in this study and representative species used in the phylogeny.

A contig containing what appears to be a complete coding sequence (3259 nt) and RdRp motifs was assembled in the *Woodsia scopulina* transcriptome and tentatively named *Rocky Mountain woodsia associated virus* (RmwPV). The predicted RdRp region (918-1921 nt) of RmvPV shares similarity to both partiti-like viruses (e.g., *Ustilaginoidea virens nonsegmented virus 2*, 26% aa identity) and the unclassified *Phytophthora infestans RNA virus 1* (42% aa identity) which has been shown to likely constitutes a novel virus family (80). The resemblance RmwPV shares with two seemingly distantly related virus groups suggest its position within the *Durnavirales* should be treated with caution (Figure 10).

The transcriptome in which RmwPV was discovered is contaminated with fungal reads (10% (Figure 2). If RmwPV was derived from fungal contaminates this could potentially explain the phylogenetic placement of RmwPV (Figure 10). The *Lycopodiella appressa* transcriptome in which LycoAV was discovered is contaminated by reads belonging to species across various land plant groups. Reads belonging to land plants comprised 35% of plant-associated reads while lycopod associated reads comprised 65% (Figure 2).

###### Partitiviridae

We detected 14 sequences that share a resemblance with members of the *Partitiviridae*. For each of these sequences, complete dsRNA1 and dsRNA2 segments were recovered. Ten sequences were found in non-flowering plants and cluster within the deltapartitiviruses. A clade within the deltpartitiviruses is known to encode a third segment comprising of a divergent dsRNA2 full-length capsid protein with unknown function (Figure 10). We identified dsRNA3 segments in related conifer associated sequences but not in those found in moss and lycophyte libraries (Figure 10). Phylogenies estimated on the coding sequences of dsRNA2 and dsRNA3 reveal essentially the same grouping which is largely consistent with the host phylogeny (SI Figure 4). We extend the known host range of the deltapartitiviruses to include liverworts, mosses, and lycophytes. The remaining sequences were found in eudicots and cluster with known plant partitiviruses (Figure 10). The white campion was judged to have contamination from ginseng and chickweed (31). However, the relatively low proportion of the library these contaminates compose (<1%) suggests that it is unlikely these species are the host of WcPV (Figure 2). There is no evidence that the other partiti-like sequences discovered are derived from contaminates.

##### Ghabrivirales

###### Chrysoviridae

We identified two partial sequences that share resemblance with members of the alphachrysoviruses denoted *Mesotaenium kramstae alphachyrso-like virus* (MkACV) and *Tree fringewort alphachyrso-like virus* (TwACV) (Figure 11). A complete RNA2 segment was recovered for MkACV which shared similarity with the p98 of various chrysoviruses (Figure 11). The MkACV RNA2 segment did not contain the “PGDGXCXXHX” motif commonly found in this protein (81). To our knowledge, these sequences represent the first chyrsoviruses in liverworts and algae. While reads belonging to fungi were found in the libraries MkACV and TwACV were assembled from, the phylogenetic positioning of the viruses suggest that they are plant-derived (Figure 2, Figure 11).

**Figure 11.**
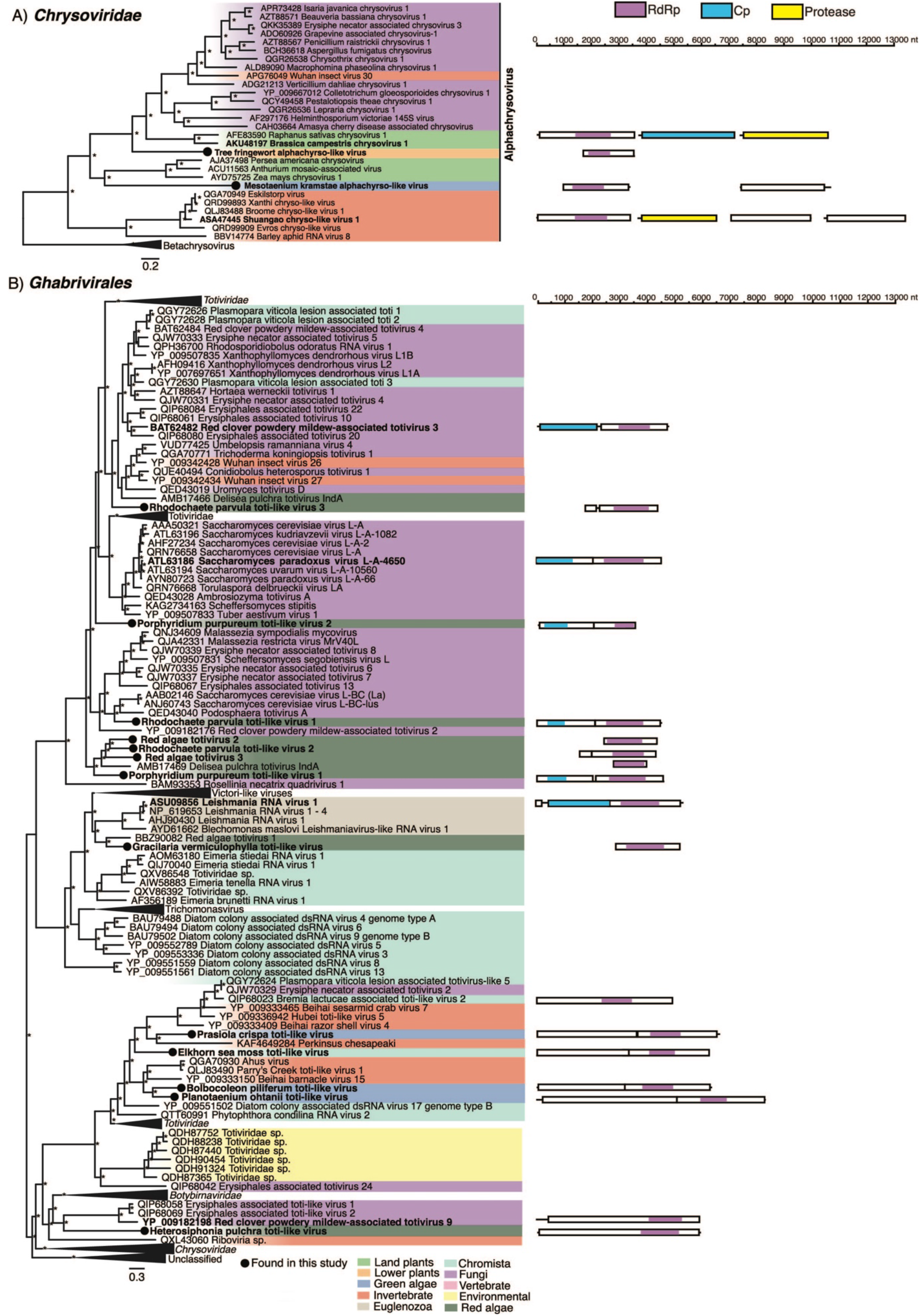
Left: Phylogenetic relationships of the viruses within the order *Ghabrivirales*. (A) A phylogeny of the *Chrysoviridae,* (B) an order level phylogeny. ML phylogenetic trees based on the replication protein show the topological position of the virus-like sequences discovered in this study (black circles) in the context of their closest relatives. See Figure 3 for the colour scheme. Green stars are used to signify sequences that have been translated using the protozoan mitochondrial genetic code. All branches are scaled to the number of amino acid substitutions per site and trees were mid-point rooted for clarity only. An asterisk indicates node support of >70% bootstrap support. Virus taxonomic names are labelled to the right. Right: Genomic organization of the virus sequences identified in this study and representative species used in the phylogeny.

###### Totiviridae

Thirteen sequences sharing similarities to toti-like viruses were discovered in eight red and green algae transcriptomes. All sequences share less than 50% amino acid identity across their coding sequence, as such we consider each a putative toti-like viruses. Among these sequences four cluster with *Delisea pulchra totivirus IndA* (AMB17469.1) to form a red alga associated clade basal to the totiviruses (Figure 11). *Gracilaria vermiculophylla toti-like virus* (GrveTV) along with *Red algae totivirus 1* (BBZ90082) form a sister group to the protozoan infecting leishmaniaviruses (Figure 11). The remaining sequences are phylogenetically positioned across the tree of toti-like viruses, commonly occupying basal positions (Figure 11).

*Prasiola crispa* is contaminated by reads from the fungi, *Candida albicans*. *Prasiola crispa toti-like virus* (PrcrTV), clusters with unclassified protist, fungi, invertebrate and algae viruses including *Elkhorn sea moss toti-like virus* (EsmTV) (Figure 2, Figure 11). The Kappaphycus alvarezii transcriptome in which EsmTV was found showed no evidence of contamination suggesting that PrcrTV may also be derived from algae (Figure 2). The *Mazzaella japonica* transcriptome in which *Red algae toti-like virus 2*:*3* (RedTV2/3) were discovered was predominantly composed of reads associated with the red algae genera Chondrus. As >99% of reads in this library belong to red algae species RedTV2 and RedTV3 have been assigned to this group. The *Porphyridium purpureum* transcriptome is highly contaminated by reads belonging to flowering plants and an unidentified cloning vector (M10197.1) (Figure 2). The phylogenetic positioning of the viruses discovered from this transcriptome (*Porphyridium purpureum toti-like virus* 1 & 2) point towards being derived from red algae rather than flowering plants (Figure 11).

### 3.4 Long-term virus-host evolutionary relationships

To examine the frequency of cross-species transmission and co-divergence among plant viruses, we estimated tanglegrams that depict pairs of rooted phylogenetic trees displaying the evolutionary relationship between a virus family and their hosts. This revealed cross-species transmission as the predominate evolutionary event predicted among all the RNA virus groups analysed (median 65%, range 46%-79%) (Figure 12). Cross-species transmission was most frequent in the *Betaflexiviridae* (79%) and the subfamily *Betarhabdovirinae* (79%). Virus-host co-divergence (median 23%, range 14%-29%) and to a lesser extent duplication (i.e., speciation) (median 4.6%, range 1.4%-24%) and extinction events (median 2.9%, range 0%-11%) were detected across plant virus families (Figure 12). Co-divergence was most frequently predicted in the *Benyviridae* and *Tymoviridae* representing 29% and 26% of events respectively. Importantly, however, the results of our co-phylogenetic analysis are undoubtedly influenced by the sample of plant viruses and will likely change as the number of plant viruses identified increases.

**Figure 12.**
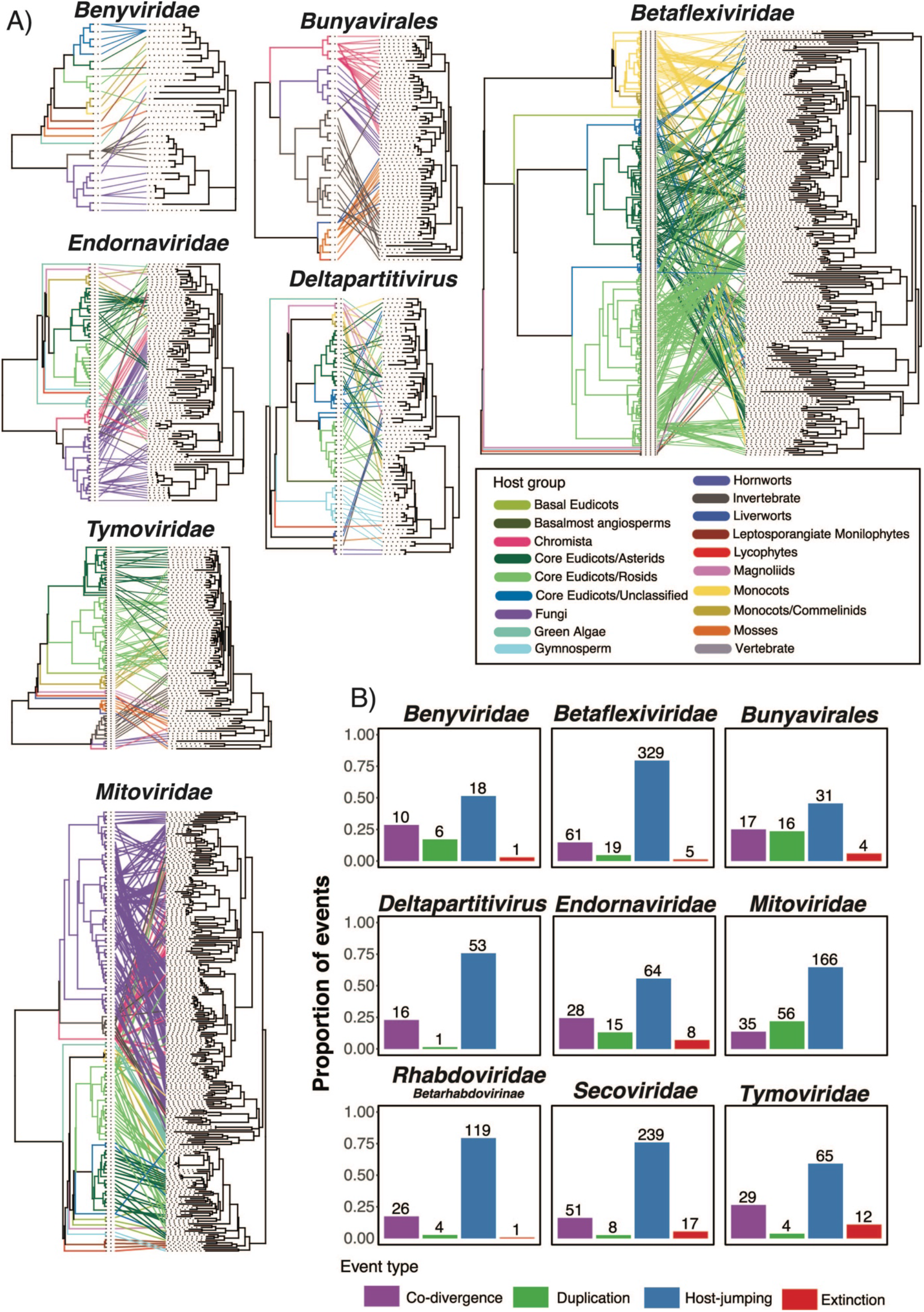
(A) Tanglegram of rooted phylogenetic trees for select virus groups and their hosts. Lines and branches are coloured to represent host clade. The cophylo function implemented in phytools (v0.7-80) was used to maximise the congruence between the host (left) and virus (right) phylogenies. Supplementary Figure 5 provides the names of the hosts and viruses along with additional tanglegrams for the *Secoviridae* and *Rhabdoviridae*. (B). Reconciliation analysis of select virus groups. Barplots illustrate the range of the proportion of possible events and are coloured by event type.

## 4. Discussion

Our ability to reconstruct the evolutionary history of plant viruses and understand the drivers of their emergence has been constrained by inadequate sampling across the enormous, extant diversity of plant species. Here, we provide a large-scale virus discovery project based on mining transcriptomes from across the entire breadth of the plant kingdom. In doing so we have identified 104 potentially novel virus species. We considerably expand upon the known host range of 13 virus families to now include lower plants and expand a further four virus families to include host associations with algae. We also find the first evidence of a movement protein with a predicted molecular weight of ∼30 kDa (herein referred to as a “30K MP”) in a virus of non-vascular plants. Collectively, this new knowledge advances our understanding of RNA virus diversity across the Archaeplastida.

### 4.1 RNA viruses are widespread across lower plant lineages

To date, viral surveys in basal plant lineages (namely ferns, bryophytes and algae) have revealed only the minimal occurrence of (+)RNA viruses (5, 17, 20, 78, 82, 83), supporting the idea that RNA viromes in angiosperms evolved as they diversified during the Cretaceous (84). However, our results potentially challenge this paradigm as we detected the first evidence of sets of (+)ssRNA viruses in lower plants and algae, implying that these groups are associated with older lineages of plants. Several of these viruses are deep branching and sit basal to angiosperm infecting viruses (e.g., LycoAV and LyBuV) in phylogenetic trees. Other viruses discovered here occupy ambiguous positions between established plant virus families (e.g., OnV) or cluster in large numbers to form novel plant-associated clades (e.g., the *Viridisbunyaviridae* in the *Bunyavirales*). Benyviruses are typically transmitted by the root-infecting plasmodiophorids *Polymyxa betae* and *Polymyxa graminis* (85, 86). The Phytomyxids (plasmodiophorids and phagomyxids) are parasites of plants, diatoms, oomycetes and brown algae and have been shown to demonstrate cross-kingdom host shifts (e.g., between angiosperms and oomycetes) (87). As such the plasmodiophorids may be a vehicle for cross-species transmission between aquatic protists and land plants (5). FeBV, a beny-like virus identified in this study, formed a clade along with *Wheat stripe mosaic virus* distinct from members of the genus *Benyvirus*. Deciphering the evolutionary history and mode of transmission for the lower plant beny-like viruses will require further studies with particular emphasis on these taxa. Interestingly, no plasmodiophorid-associated reads were detected in any of the libraries from which we assembled a beny-like virus. LjBV and WasBV appear distantly related to the benyviruses. These viruses group with a suite of unclassified viruses assembled from a soil metatranscriptome study suggesting that, like the benyviruses, this larger group of unclassified viruses may involve soil-borne parasites like the plasmodiophorids (88). Our detection of tymovirid-like sequences in the lycophytes, bryophytes and brown algae dramatically expands the known host range of the *Tymovirales*. Several of these viruses were similar to unclassified Riboviria species assembled from a recent survey of common wild oat soil rhizosphere and detritosphere (88) (Figure 4). The metatranscriptome of the sequenced soil samples from the common wild oat study was largely composed of Viridiplantae, fungi, Amoebozoa, protists, nematodes, and other eukaryotes. As such, using phylogenetic clusters to infer host associations of our viruses remains challenging. Indeed, these viruses may result from contamination from other eukaryotes (e.g., fungi or invertebrates) although we found no consistent evidence among these viruses (Figure 2). Assuming these viruses are plant-associated, their phylogenetic pattern suggests that they may have resulted from cross-kingdom transmission events that frequent the Alsuviricetes.

The partial deltaflexi-like virus we detected in *P. agnata* (PaADV) is particularly noteworthy. The deltaflexiviruses are only known to infect fungi, although no fungi associated reads were found in the *P. agnata* metatranscriptome (Figure 2). The mycovirus families *Delta*- and *Gammaflexiviridae* are thought to have been derived from the plant alpha- and betaflexivirids through cross-species transmission (5, 89)). As such PaAGV could potentially represent an intermediate between the plant and fungi flexiviruses or perhaps a more recent fungus to plant transmission. As only a fragment of the polymerase gene was assembled for this virus future work should confirm the presence of PaAGV and its phylogenetic position.

### 4.2 The extension of the *Mitoviridae* to a lycophyte host

Through the analysis of mitoviruses-like, non-retroviral endogenous RNA viral elements (NERVEs), it was argued that the origin of plant mitovirus NERVEs was a single horizontal transfer from a fungal mitovirus before the origin of vascular plants in the early Silurian, ∼400 MYA (90). Evidence of contemporary mitoviruses in flowering plants and a fern have challenged this view, suggesting that a lineage of plant rather than fungal mitoviruses are the immediate ancestors of plant mitovirus NERVEs (16). Indeed, plant-to-fungus transmission would eliminate code conflicts between fungi and plant mitochondrial genetic codes (76). Herein, we demonstrate the existence of a lower plant-associated sister clade to the angiosperm mitoviruses and NERVEs. This clade includes a clubmoss associated mitovirus, the most primitive plant mitovirus sequence to date. This finding aligns with the estimation of the origin of plant mitovirus NERVEs occurring as early as the evolution of the clubmoss (90). The recent finding of mitoviruses in green algae – including BopiMV in this study – highlight the broad host range of mitoviruses (78, 83). The phylogenetic position of these viruses and the absence of NERVEs from these groups suggest that they are not the ancestors of land plant mitoviruses and NERVEs.

### 4.4 Establishment of a new virus family in the *Bunyavirales*: *Viridisbunyaviridae*

We identified 16 bunya-like viruses assembled from six non-vascular plant libraries including liverwort, moss, and lycophyte species. These viruses form a novel clade within the *Bunyavirales* and represent the first viruses in this order to be associated with lower plants. This clade likely represents a novel virus family which we have tentatively named the *Viridisbunyaviridae.* Several libraries contained up to five distinct viruses (each sharing <70% nucleotide identity). Virus co-infections are frequently observed in plants and have been reported in the closest relatives of these viruses, the *Deltamycobunyaviridae* (91, 92). As with previous studies we were only able to recover the bunyavirus L segment (92, 93). Further studies are needed to recover the missing small and medium-sized segments and to confirm the presence of mixed infections in plants.

### 4.5 Discovery of the first 30 kDa movement protein in non-vascular plants

Through the discovery of lower plant-associated viruses, we have gained insights into how the genome structure and composition of contemporary flowering plants viruses have evolved. The detection of secovirid-like sequences in bryophytes and ferns represents the first occurrence of plant secoviruses outside of angiosperms and the first evidence of a 30K MP homolog in non-vascular plants. These proteins aid the cell-to-cell movement of viruses in plants. For example, the MP of *Cucumber mosaic virus* increases the size exclusion limit of plasmodesmata allowing virus particles to pass through cell walls (94). To date, homologs of 30K MP have only been detected in plant viruses infecting angiosperms to the lycophytes (17, 95). Further work is needed to confirm the presence and function of 30K MPs in viruses infecting the bryophytes and other lower plants.

### 4.6 Detection of Deltaparitivirus dsRNA3 segments in gymnosperms but not in non-vascular plants

Our discovery of six tri-segmented deltapartitivirus species provides insights into the evolution of the deltapartitivirus dsRNA3 segment. dsRNA3 segments have been found in several alpha- and deltapartitiviruses infecting flowering plants (96–99). These segments typically encode seemingly full-length capsid protein or in the case of alphapartitivirus *Rosellinia necatrix partitivirus 2*, a truncated version of the RdRp which may serve as an interfering RNA (100). There is some debate as to the source of dsRNA3 segments, particularly whether they are satellite viruses that co-opt the RdRp of the co-infecting helper viruses or that the additional segment is a result of coinfection of two different plant partitiviruses and the second RdRp-encoding segment is lost after the initial infection (101). For the first time, we find dsRNA3 segments in conifer associated viruses but not in those found in lower plants including bryophytes and lycophytes. The absence of dsRNA3 in non-vascular plants means that it is possible that this segment evolved after the divergence of vascular and non-vascular plants in the Silurian period (102). It is possible that dsRNA3 segments exist for the non-vascular plant infecting deltapartitiviruses but was not detected due to the large degree of divergence between this segment and reference sequences (including those found in this study). However, the dsRNA1 and dsRNA2 segments of the putative lower plant deltapartitiviruses shared >50% aa identity with the tri-segmented deltapartitiviruses - well above the detection limit for tools such as Diamond BlastX (37). dsRNA3 segments typically appear no more divergent than dsRNA2 segments therefore it is unlikely that we would be able to detect both the dsRNA1 and dsRNA2 segments without detecting dsRNA3. Further work is needed to confirm the presence of deltapartitivirus dsRNA3 segments.

### 4.7 Discovery of an unsegmented varicosavirus-like viruses in ferns and liverworts

Finally, the recently discovered gymnosperm varicose-like *Pinus flexilis virus 1* in the family *Rhabdoviridae* contains an unsegmented genome organisation that differs from the typical bi-segmented structure of the varicosaviruses (25, 103). We find the bi-segmented structure in varicosavirus-like viruses for the first time in ferns and liverworts (TfVV and MgVV) which predate the gymnosperms.

### 4.8 Caveats

Importantly, the data generated under the 1KP were not explicitly created for virus discovery, such that there are important caveats associated with the methods and metatranscriptomic data mined for virus contigs. For instance, as axenic cultures are not a viable option in most instances, the 1KP samples are commonly contaminated by nucleic acids belonging to bacterial, fungal, and insect species. We addressed this by using a combination of host/virus abundance measurements and phylogenetic analyses to improve the accuracy of virus-host assignments. For most of the viruses described, phylogenetic placement within plant infecting virus families strongly supports their association with plants. However, several of the viruses found in algae and lower plants were associated with lineages known to infect invertebrates and fungi or unclassified viruses recovered from environmental samples. The association between the viruses of lower plants and algae with that of fungi and invertebrate viruses may reflect the absence of algal and lower plant viruses in reference sequence databases. Experimental confirmation is needed to formally assign the viruses discovered in this study to their hosts.

The average sequencing depth of the 1KP libraries was 1.99 gigabases of sequence per sample (range 1.3-3.0), lower than many other virus discovery studies (6, 104, 105). Sequencing depth has been shown to correlate with the ability to detect viruses present at low abundance (106, 107). Further, a large proportion of the virus transcripts detected were from viruses whose full-length genomic or subgenomic mRNAs were polyadenylated at the 3′ end (SI Table 4, Figure 1). Although this was anticipated (i.e. the libraries generated by the 1KP initiative were prepared from polyA+ RNA), it limited the detection of non-polyadenylated viruses (e.g., dsRNA, dsDNA) and may have contributed to the lack of phycodnavirus sequences detected in algae (107).

To reduce the computational burden of assembly, we attempted to remove host-associated reads before contig assembly by mapping them to the host scaffolds provided by the 1KP initiative. While this step reduces the occurrence of false-positive virus detection it also risks removing virus reads, particularly reverse-transcribing plant viruses (108). While we frequently detected transcripts associated with the reverse-transcribing family *Caulimoviridae,* no members of the *Metaviridae* or *Pseudoviridae* were detected.

## Acknowledgements

We thank Richard Miller for computational support and Justine Charon for advice on alignments. This work would not have been possible without the data generously provided by the One Thousand Plant Transcriptomes Initiative.

## Supplementary Information

**Supplementary Figure 1.** (A) Phylogram of the triple gene block (TGB) protein 1. ML phylogenetic trees show the topological position of the newly discovered TGB sequence in the tomato fern (black circle) in the context of the closest relatives. (B) Phylogram of the Tymoviridae virus coat proteins (CP). ML phylogenetic trees show the topological position of the newly discovered CP sequences in (black circle) in the context of the closest relatives. Branches are highlighted to represent virus taxonomy (Maculavirus = green, Marafivirus = orange, Tymovirus = red and unclassified = grey). For each colour, a lighter shade signifies that this virus is related to but has not formally been assigned to this genus. (C) Phylogram of the *Oxera neriifolia associated virus* coat protein (CP). ML phylogenetic trees show the topological position of the newly discovered CP sequence (black circle) in the context of the closest relatives. For all trees, branches are scaled to the number of amino acid substitutions per site and trees were mid-point rooted for clarity only. Numbers at the nodes indicate bootstrap support over 70% (1000 replicates).

**Supplementary Figure 2.** Multiple sequence alignment conserved amino acid motifs in RNA-dependent RNA polymerase (RdRp) regions of the mitoviruses discovered in this study along with reference mitoviruses. The bar above each residue is green if 100% of residues in that column are identical, green-brown if they are 30%-99%, and red if under 30%. The numbers under each section correspond to regions containing motifs identified in (72).

**Supplementary Figure 3.** (A) Phylogenetic relationships of the viruses identified within the virus families *Potyviridae and Tombusviridae*. ML phylogenetic trees based upon alignments of the amino acid sequences of the RdRp protein show the topological position of discovered virus-like sequences (black circles) from this study in the context of their closest relatives. See Figure 3 for the colour scheme. All branches are scaled to the number of amino acid substitutions per site and trees were mid-point rooted for clarity only. An asterisk indicates node support of >70% bootstrap support.

**Supplementary Figure 4.** Phylogram of the deltapartitii-like virus (A) coat protein/RNA2 (CP) and (B) RNA3/coat protein 2. ML phylogenetic trees show the topological position of the newly discovered CP sequences in (black circle) in the context of the closest relatives. All branches are scaled to the number of amino acid substitutions per site and trees were mid-point rooted for clarity only. Numbers at the nodes indicate bootstrap support over 70% (1000 replicates).

**Supplementary Figure 5.** Tanglegram of rooted phylogenetic trees for select virus families and their hosts. Lines and branches are coloured to represent host clade. The cophylo function implemented in phytools (v0.7-80) was used to maximise the congruence between the host (left) and virus (right) phylogenies.

**Supplementary Table 1.** Clade assignment for all One Thousand Plant Transcriptomes Initiative (1KP) species for which a virus was detected.

**Supplementary Table 2.** Summary information for each One Thousand Plant Transcriptomes Initiative (1KP) libraries analysed.

**Supplementary Table 3.** Proportion of transcripts and abundance assigned to each plant virus family.

**Supplementary Table 4.** Summary table of the viruses discovered in this study

**Supplementary Table 5.** Genome annotation information underlying the annotation graphs

