## Supplementary figures and images for "Transcriptome Mining Reveals a Spectrum of RNA Viruses in Primitive Plants"

### Supplementary Figure 1

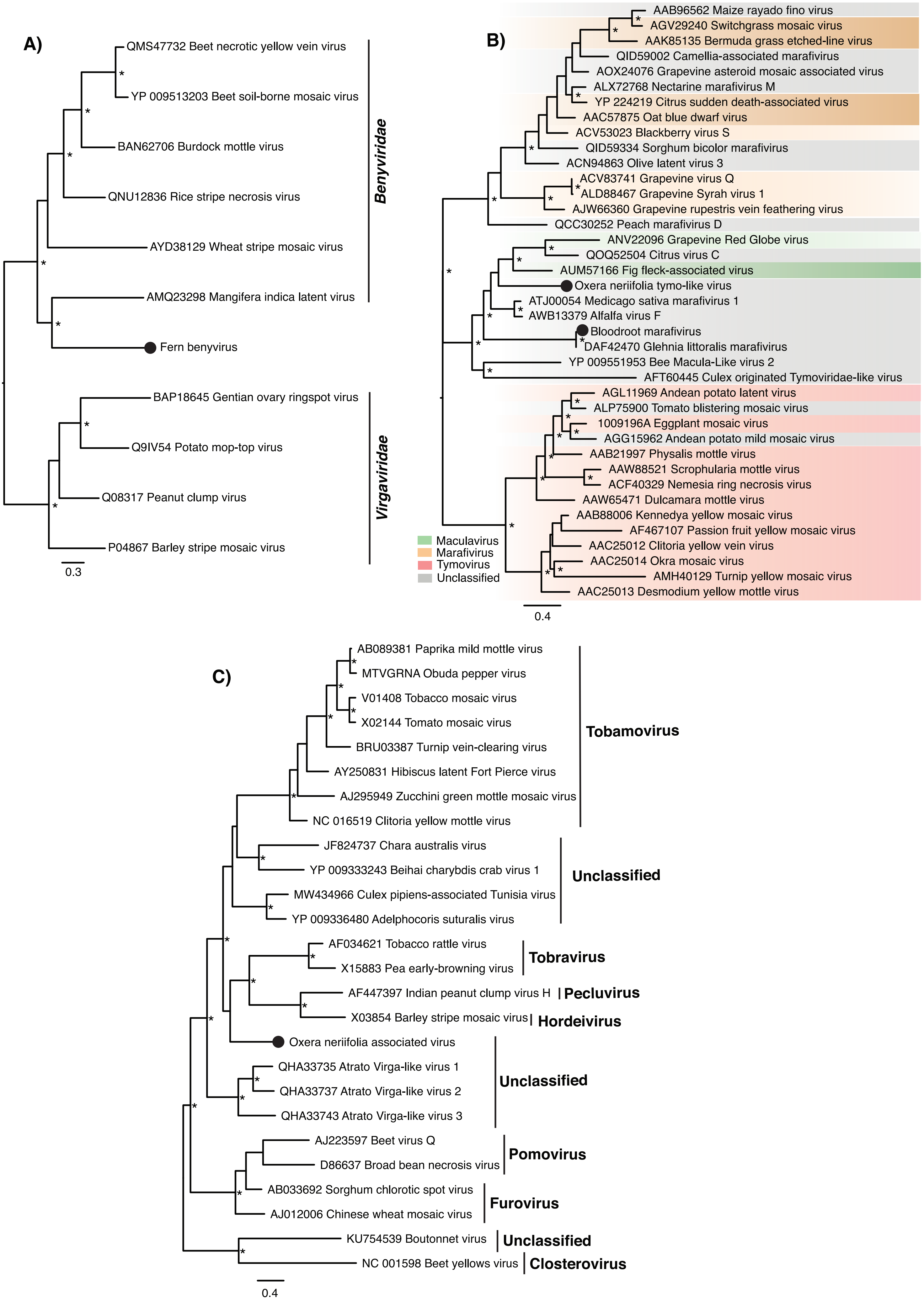

### Supplementary Figure 2

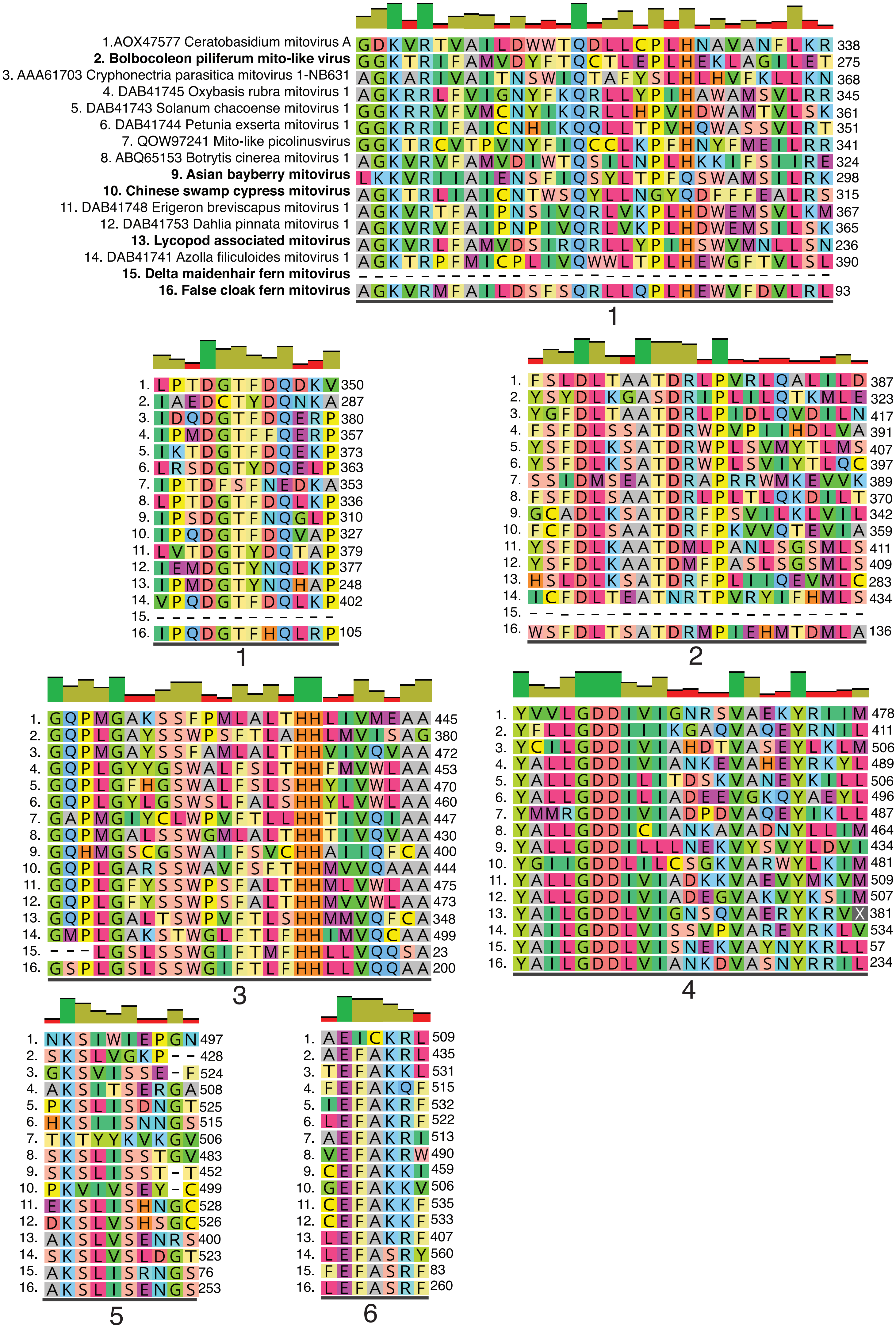

### Supplementary Figure 3

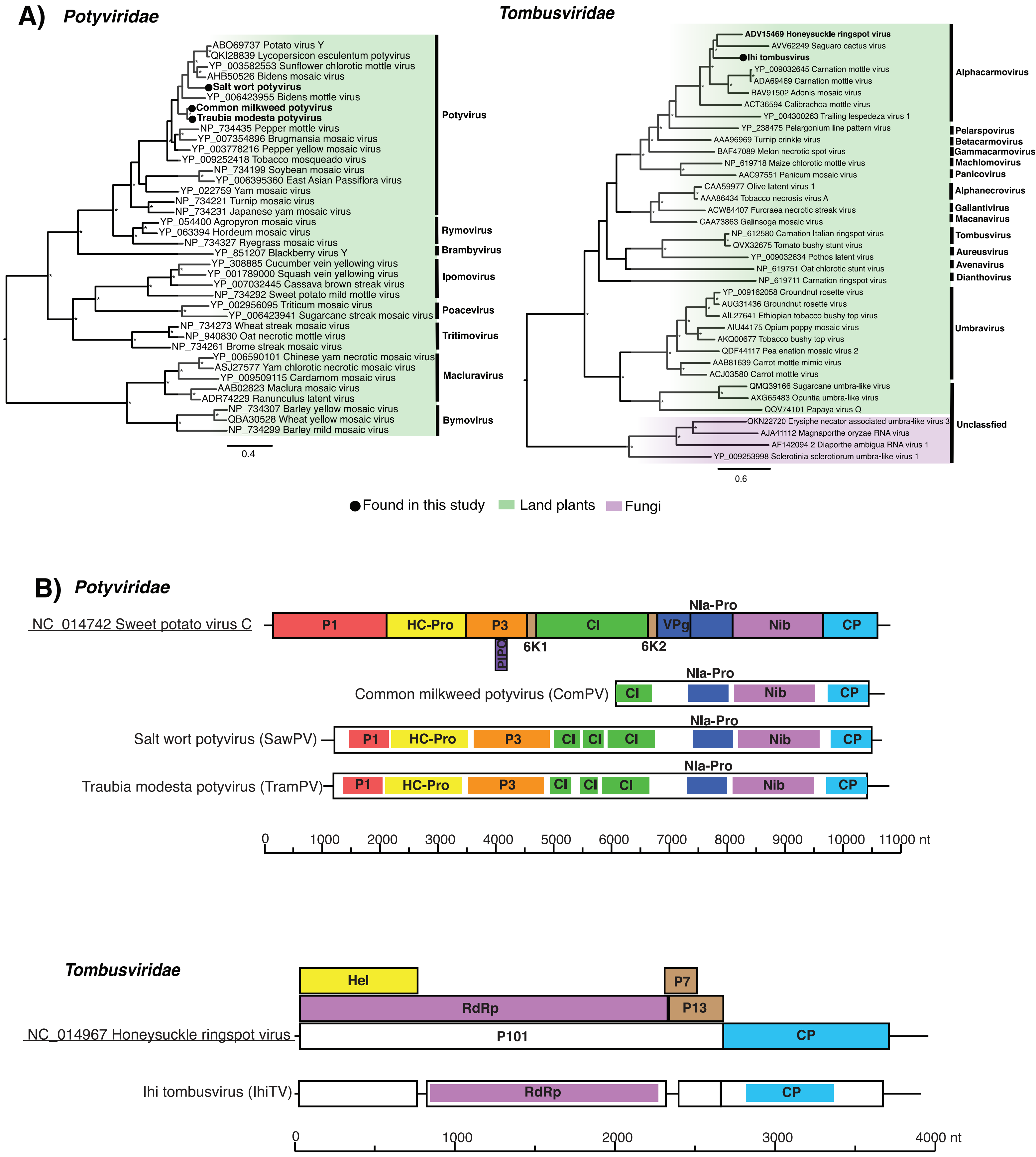

### Supplementary Figure 4

A) **CP1/RNA2**

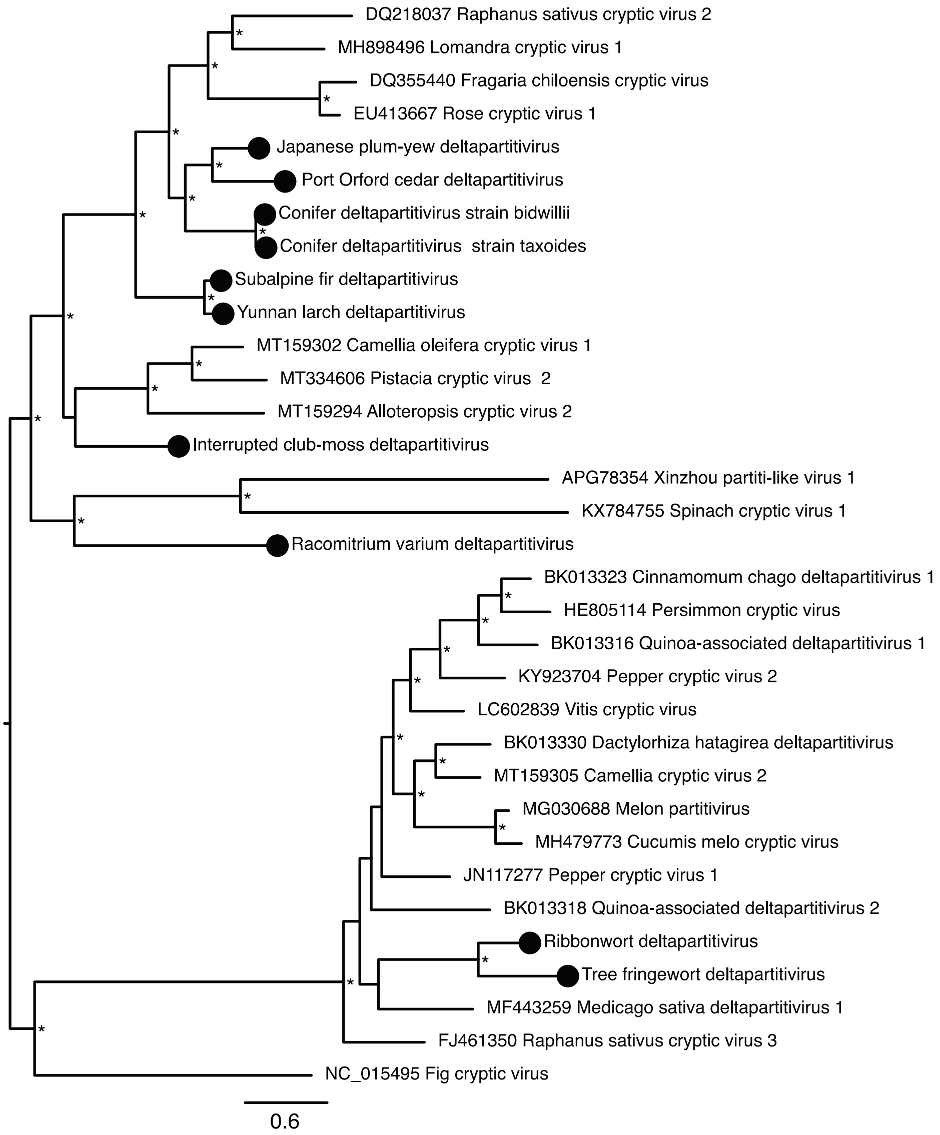


B) **CP2/RNA3**

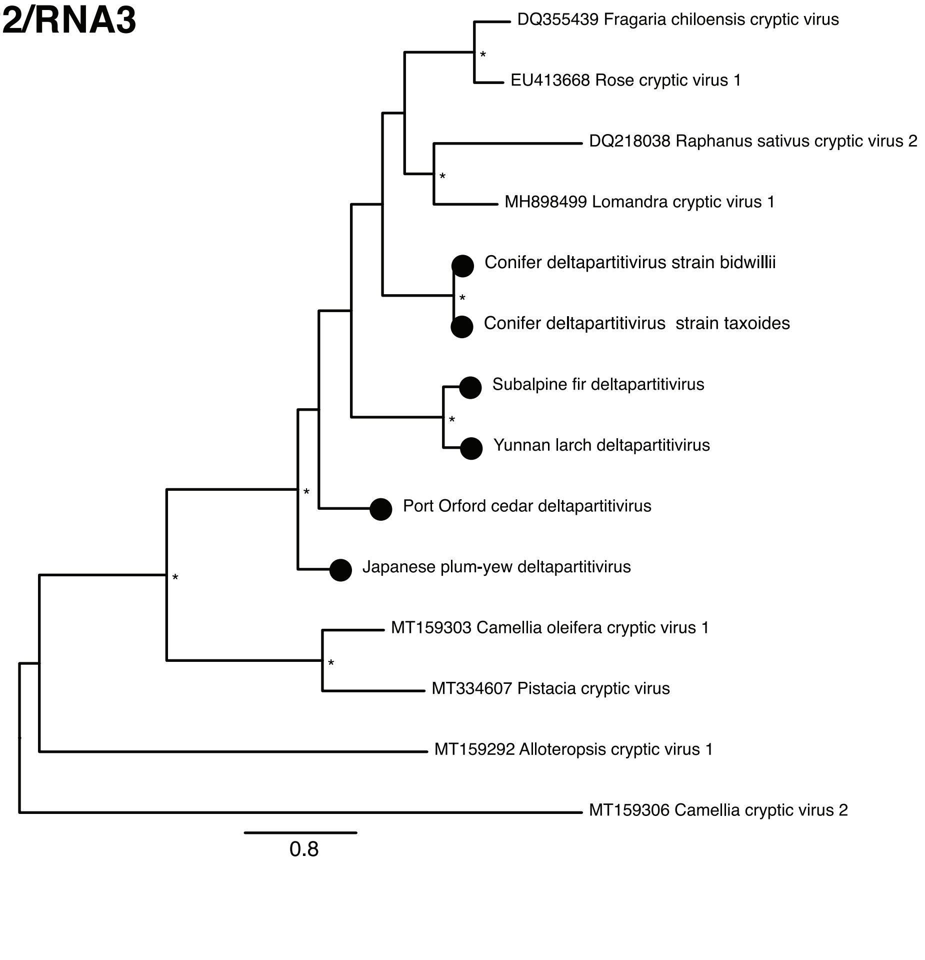
