## Supplementary Figure 5 for "Transcriptome Mining Reveals a Spectrum of RNA Viruses in Primitive Plants"

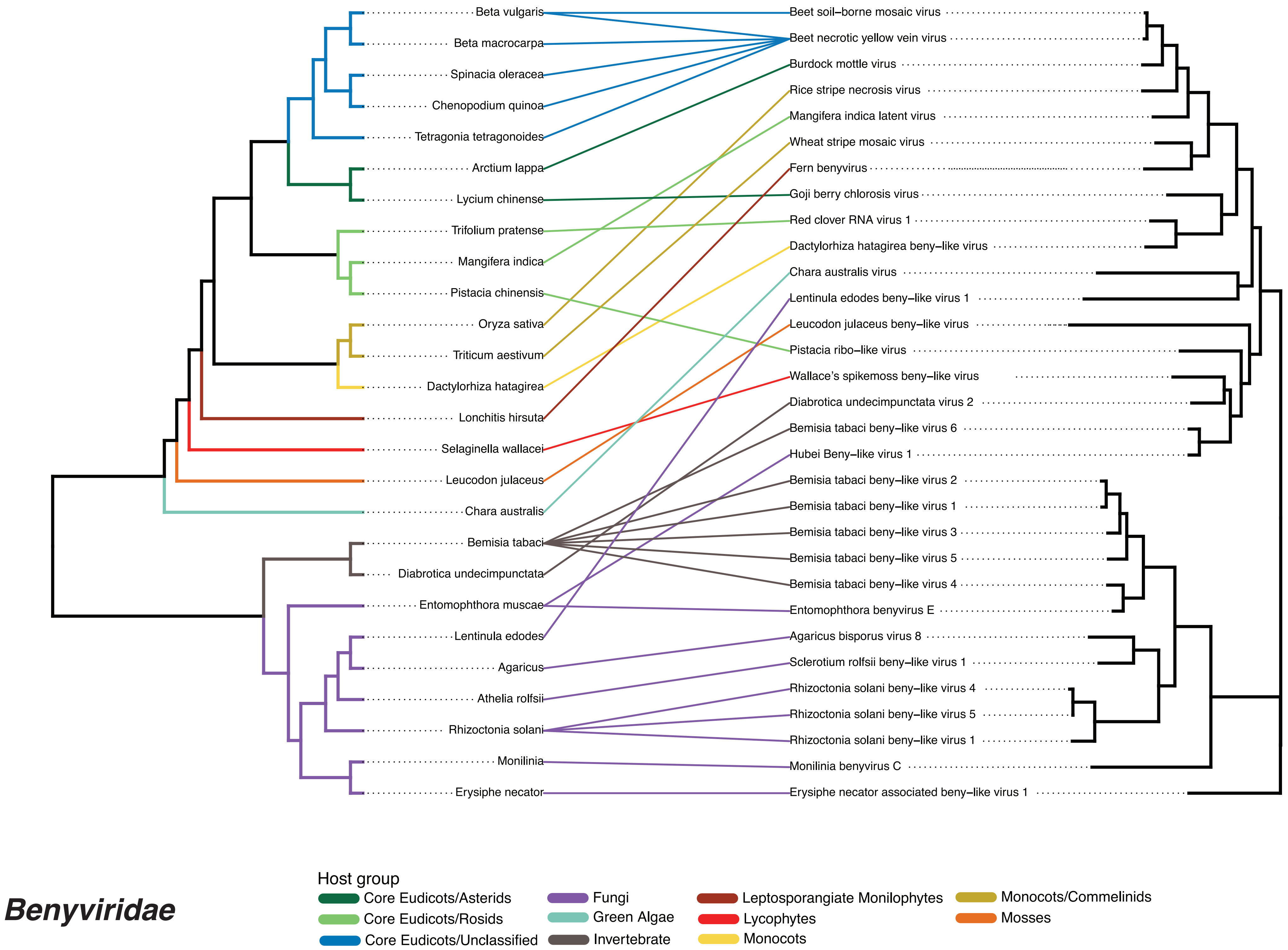

Beta vulgaris
Beta macrocarpa
Spinacia oleracea
Chenopodium quinoa
Tetragonia tetragonoides
Arctium lappa
Lycium chinense
Trifolium pratense
Mangifera indica
Pistacia chinensis
Oryza sativa
Triticum aestivum
Dactylorhiza hatagirea
Lonchitis hirsuta
Selaginella wallacei
Leucodon julaceus
Chara australis
Bemisia tabaci
Diabrotica undecimpunctata
Entomophthora muscae
Lentinula edodes
Agaricus
Athelia rolfsii
Rhizoctonia solani
Monilinia
Erysiphe necator
Beet soil–borne mosaic virus
Beet necrotic yellow vein virus
Burdock mottle virus
Rice stripe necrosis virus
Mangifera indica latent virus
Wheat stripe mosaic virus
Fern benyvirus
Goji berry chlorosis virus
Red clover RNA virus 1
Dactylorhiza hatagirea beny–like virus
Chara australis virus
Lentinula edodes beny–like virus 1
Leucodon julaceus beny–like virus
Pistacia ribo–like virus
Wallace's spikemoss beny–like virus
Diabrotica undecimpunctata virus 2
Bemisia tabaci beny–like virus 6
Hubei Beny–like virus 1
Bemisia tabaci beny–like virus 2
Bemisia tabaci beny–like virus 1
Bemisia tabaci beny–like virus 3
Bemisia tabaci beny–like virus 5
Bemisia tabaci beny–like virus 4
Entomophthora benyvirus E
Agaricus bisporus virus 8
Sclerotium rolfsii beny–like virus 1
Rhizoctonia solani beny–like virus 4
Rhizoctonia solani beny–like virus 5
Rhizoctonia solani beny–like virus 1
Monilinia benyvirus C
Erysiphe necator associated beny–like virus 1
Host group
Core Eudicots/Asterids
Core Eudicots/Rosids
Core Eudicots/Unclassified
Fungi
Green Algae
Invertebrate
Leptosporangiate Monilophytes
Lycophytes
Monocots
Monocots/Commelinids
Mosses
Benyviridae

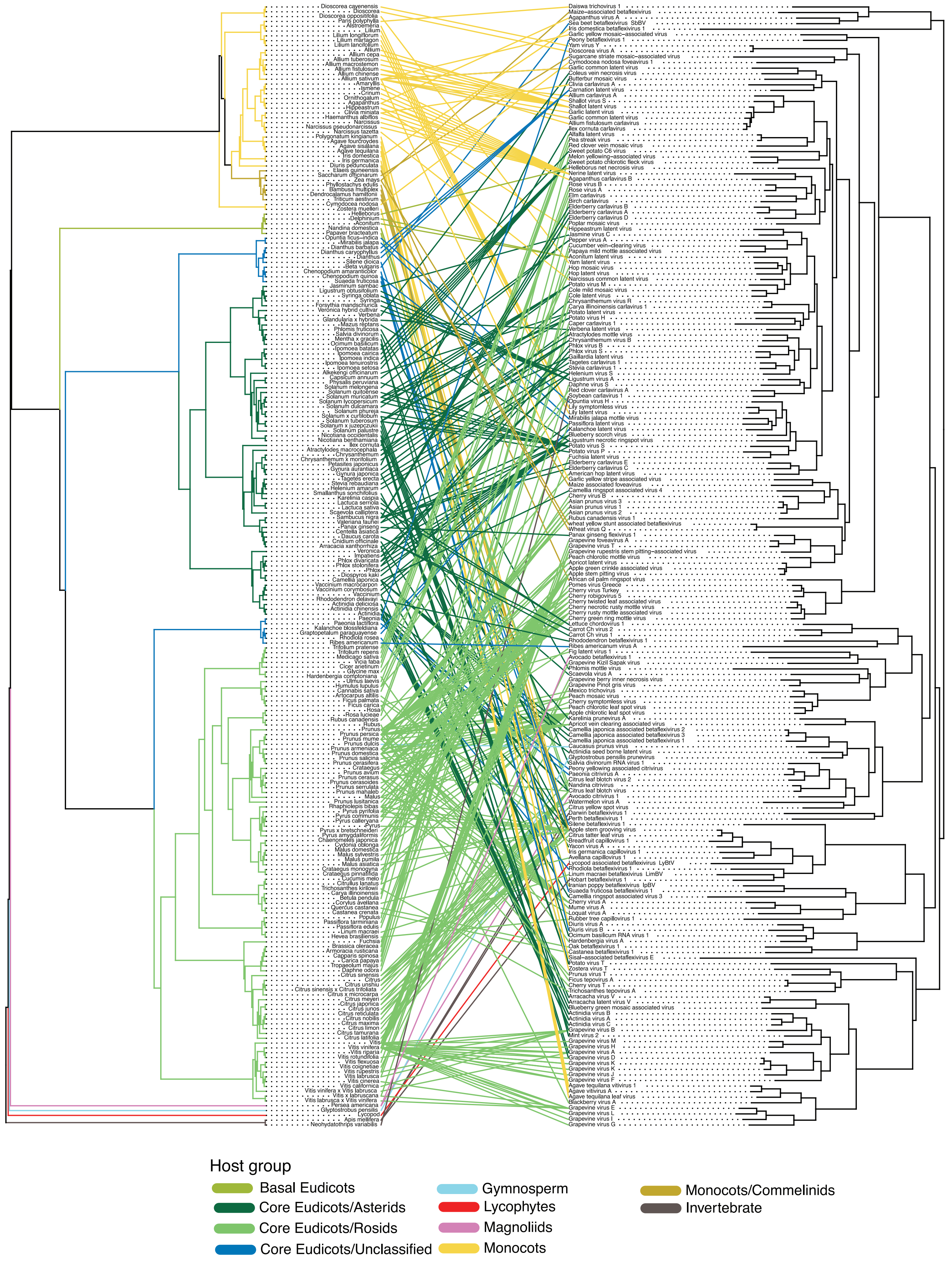


***Betaflexiviridae***

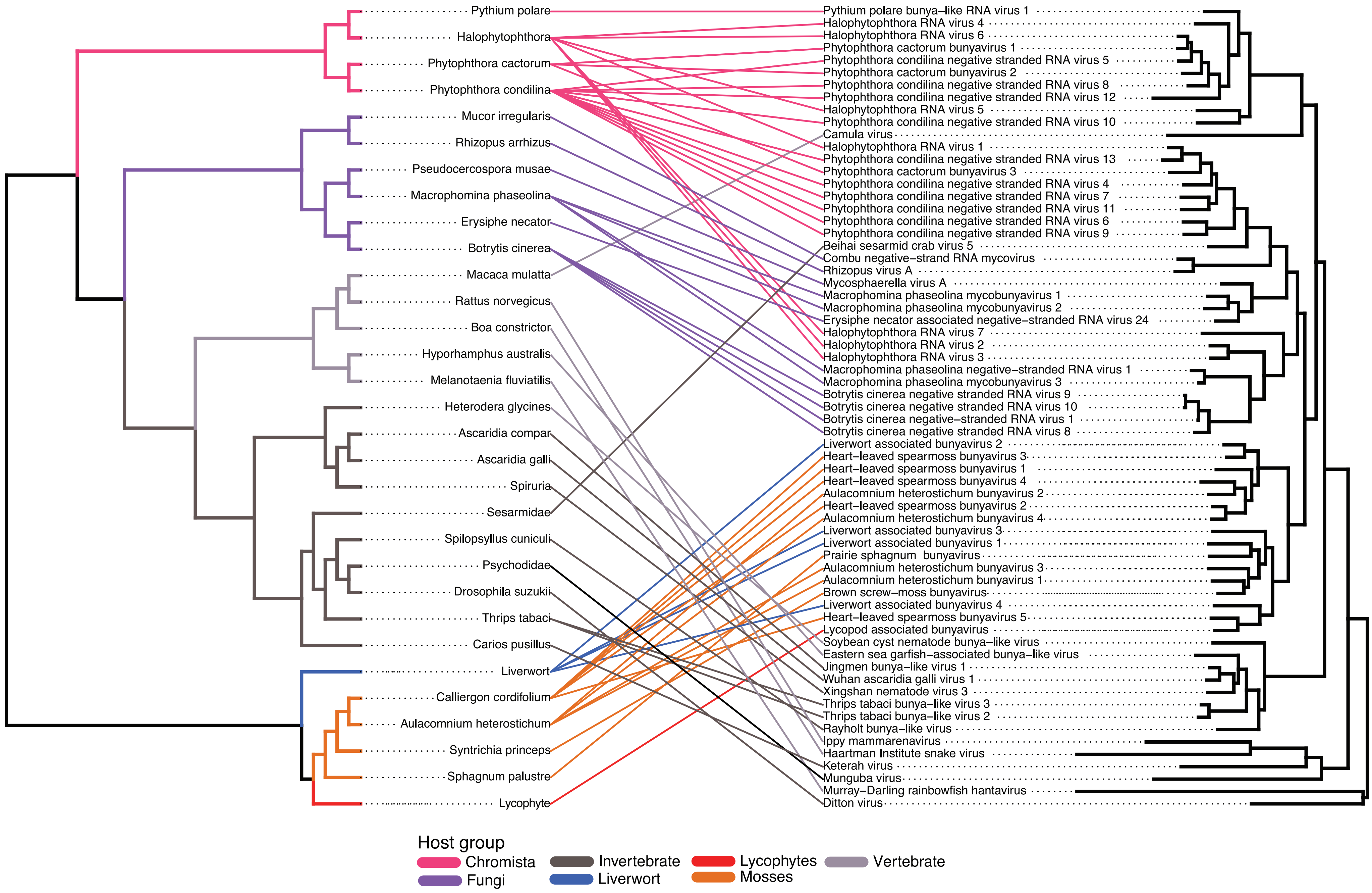


***Bunyavirales/Viridisbunyaviridae***

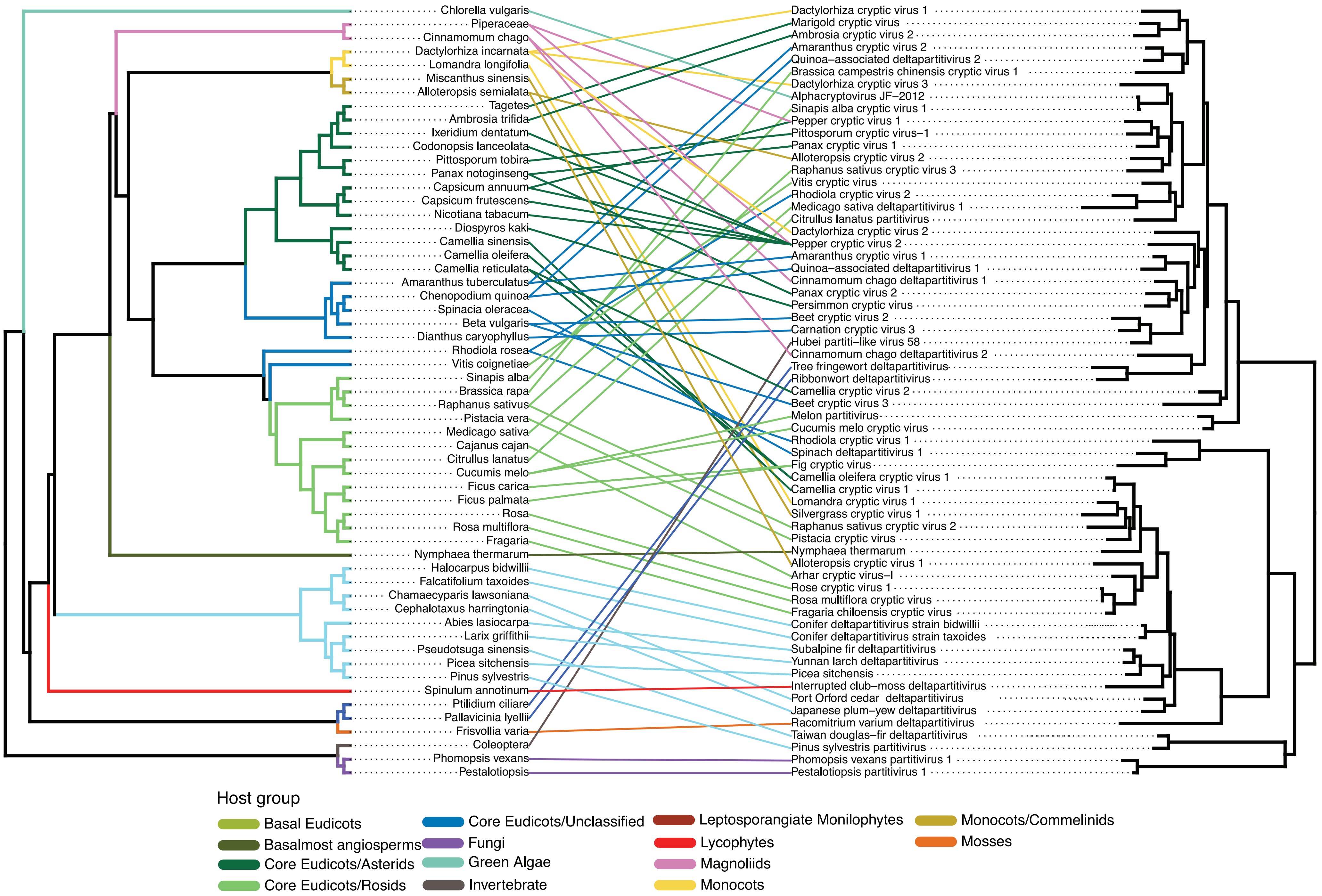


***Deltapartitivirus***

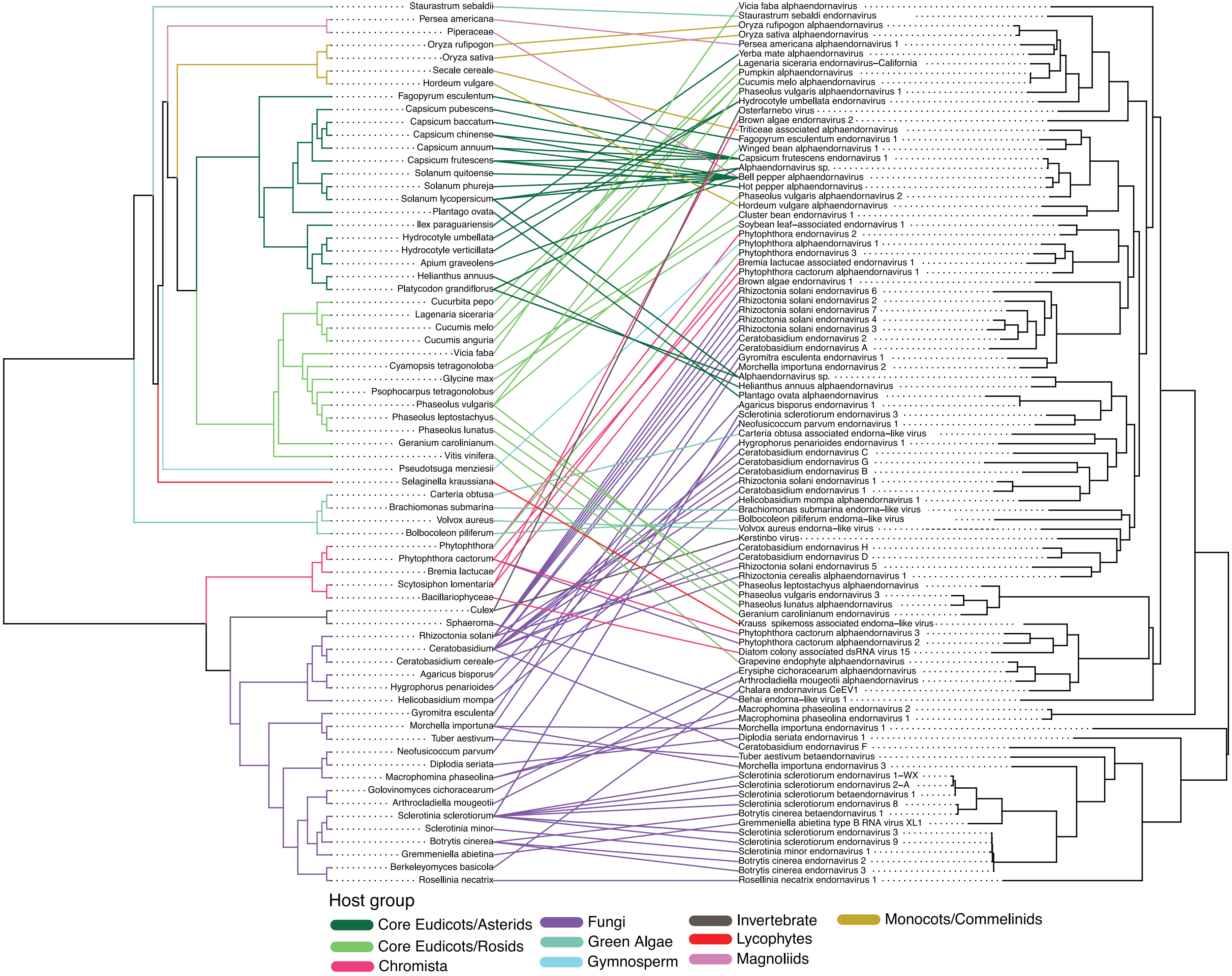


***Endornaviridae***

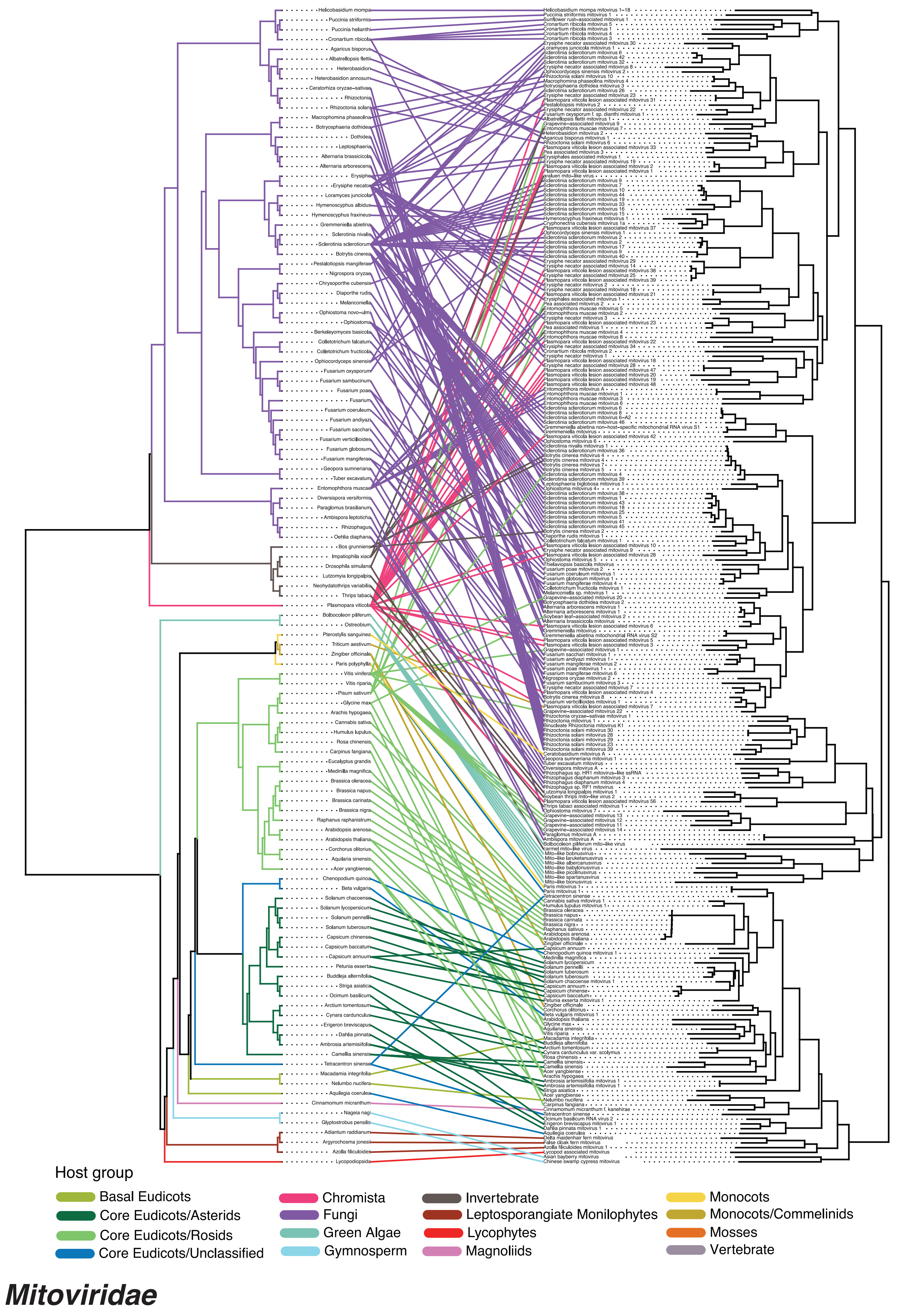

Host group
Basal Eudicots
Core Eudicots/Asterids
Core Eudicots/Rosids
Core Eudicots/Unclassified
Chromista
Fungi
Green Algae
Gymnosperm
Invertebrate
Leptosporangiate Monilophytes
Lycophytes
Magnoliids
Monocots
Monocots/Commelinids
Mosses
Vertebrate
Mitoviridae

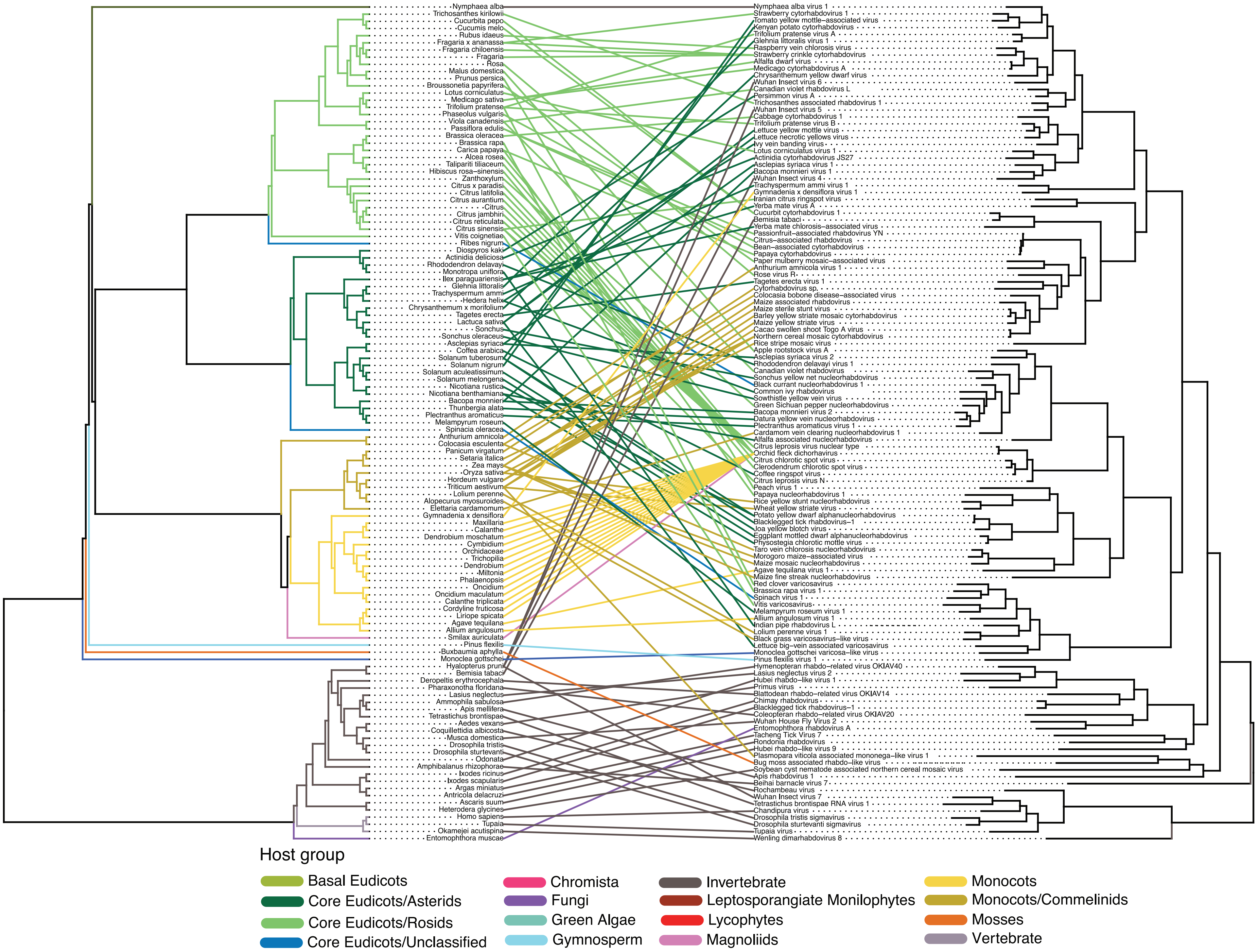


***Rhabdoviridae/Betarhabdovirinae***

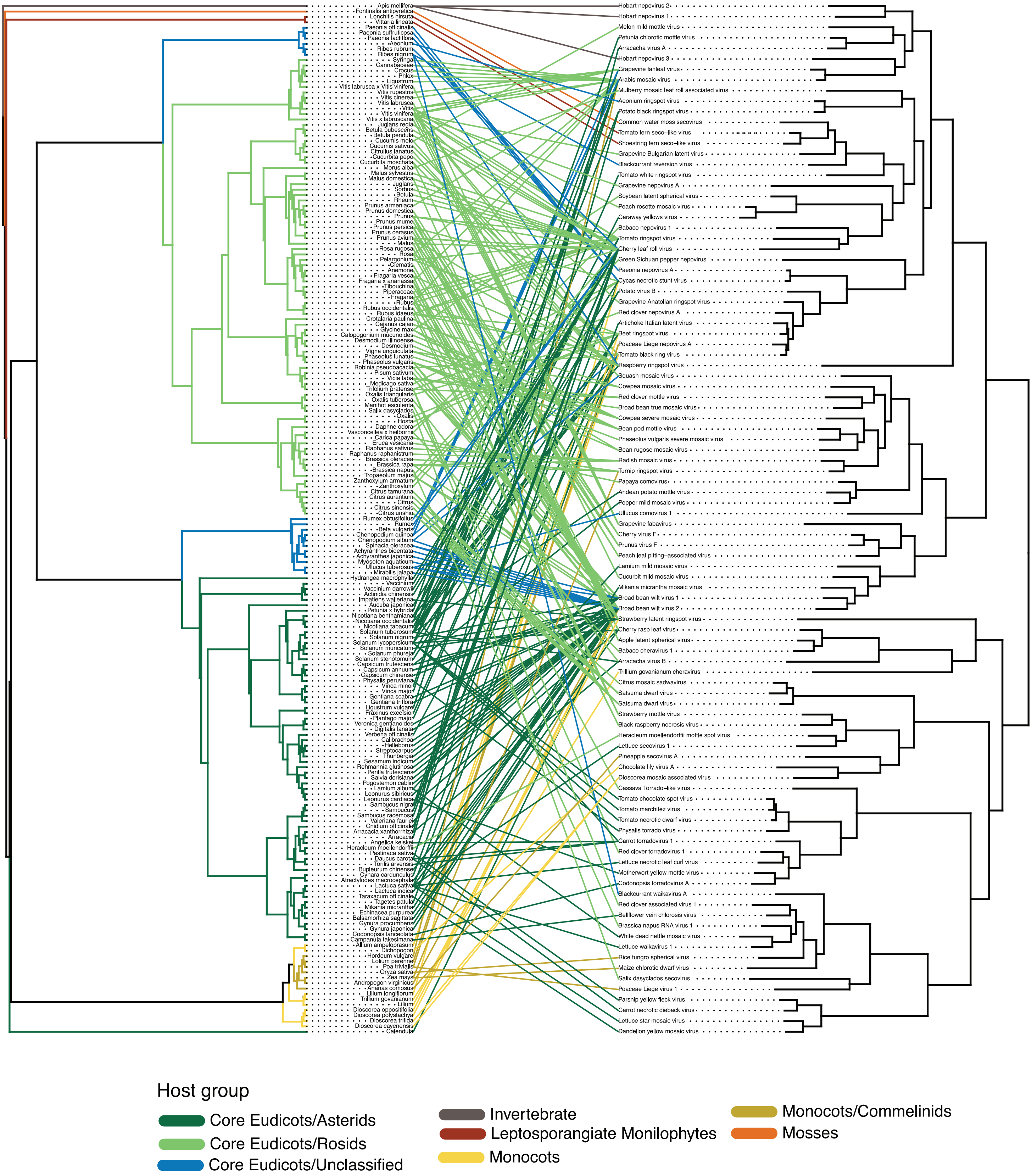


***Secoviridae***

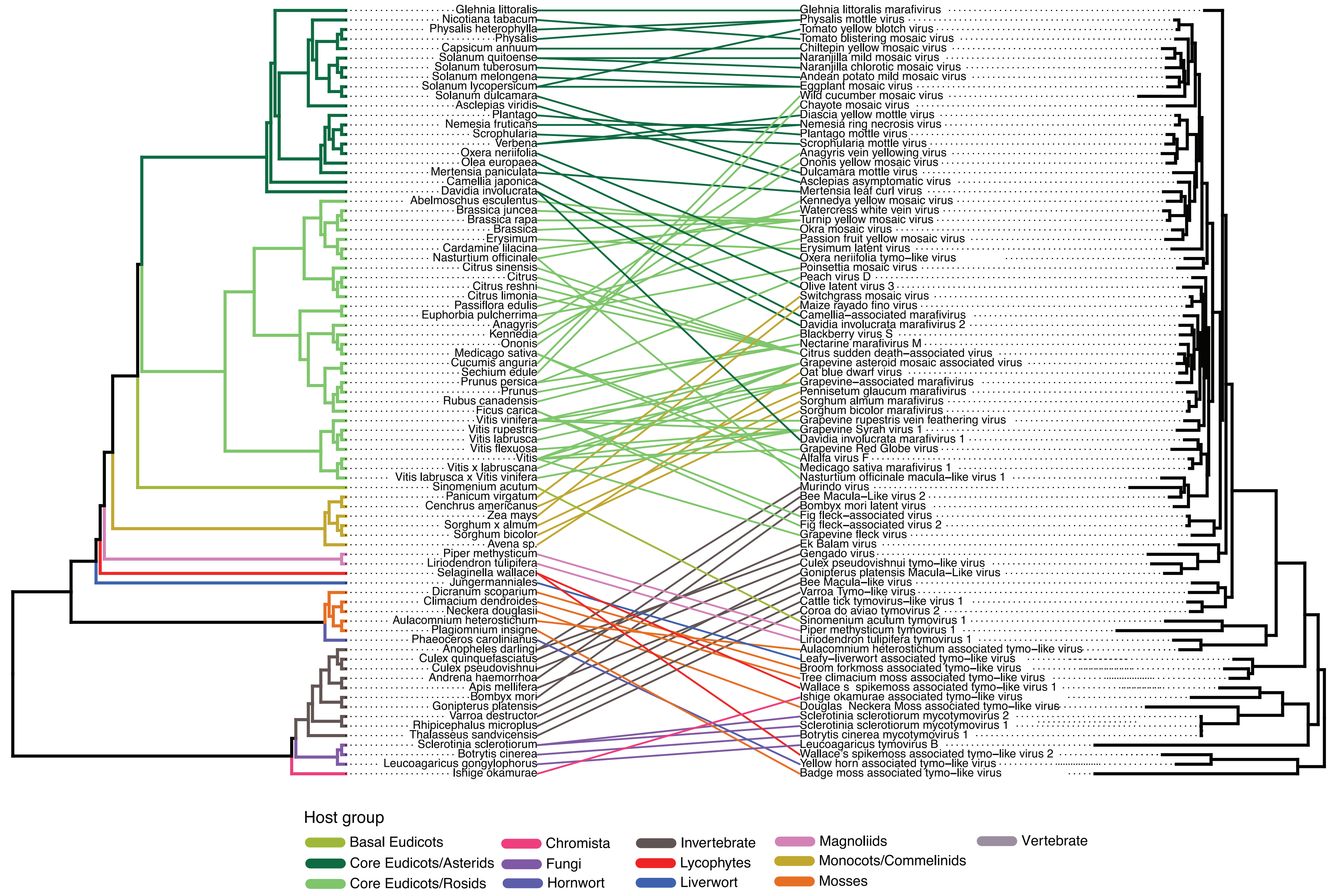


***Tymoviridae***
