## Supplementary Table 3 for "Transcriptome Mining Reveals a Spectrum of RNA Viruses in Primitive Plants"

**Supplementary Table 3: Percentage of transcripts and abundance assigned to each plant virus family**

| Virus family | Percentage of total transcripts | Proportion of total virus transcript abundance |
|---|---|---|
| *Alphaflexiviridae* | 7% | 40% |
| *Amalgaviridae* | 1% | 0.001% |
| *Aspiviridae* | 0.04% | 0.001% |
| *Benyviridae* | 0.3% | 0.003% |
| *Betaflexiviridae* | 20% | 38% |
| *Bromoviridae* | 1% | 0.1% |
| *Caulimoviridae* | 21% | 0.1% |
| *Chrysoviridae* | 0.3% | 0.0004% |
| *Closteroviridae* | 1% | 0.002% |
| *Endornaviridae* | 0.3% | 0.001% |
| *Geminiviridae* | 2% | 0.01% |
| *Luteoviridae* | 1% | 0.003% |
| *Marnaviridae* | 0.5% | 0.01% |
| *Partitiviridae* | 7% | 0.1% |
| *Potyviridae* | 13% | 15% |
| *Rhabdoviridae* | 6% | 0.009% |
| *Secoviridae* | 11% | 7% |
| *Tombusviridae* | 0.2% | 0.01% |
| *Tospoviridae* | 3% | 0.03% |
| *Tymoviridae* | 2% | 0.1% |
| *Unclassified* | 2% | 0.007% |
| *Virgaviridae* | 0.4% | 0.002% |
